# 1-Deoxysphingolipids Tempt Autophagy Resulting in Lysosomal Lipid Substrate Accumulation: Tracing the Impact of 1-Deoxysphingolipids on Ultra-Structural Level using a Novel Click-Chemistry Detection

**DOI:** 10.1101/2021.01.21.427595

**Authors:** Christian Lamberz, Marina Hesse, Gregor Kirfel

## Abstract

Sphingolipids (SLs) are pivotal components of biological membranes essentially contributing to their physiological functions. 1-deoxysphingolipids (deoxySLs), an atypical cytotoxic acting sub-class of SLs, is relevant for cellular energy homeostasis and is known to be connected to neurodegenerative disorders including diabetic neuropathy and hereditary sensory neuropathy type 1 (HSAN1). High levels of deoxySLs affect lipid membrane integrity in artificial liposomes. Accordingly, recent reports questioned the impact of deoxySLs on physiological lipid membrane and organelle functions leading to impaired cellular energy homeostasis.

However, DeoxySL-related structural effects on cell membranes resulting in organelle dysfunction are still obscure. To illuminate disease-relevant sub-cellular targets of deoxySLs, we traced alkyne-containing 1-deoxysphinganine (alkyne-DOXSA) and resulting metabolites on ultra-structural level using a new labeling approach for electron microscopy (EM) termed “Golden-Click-Method” (GCM). To complement high-resolution analysis with membrane dynamics, selected intracellular compartments were traced using fluorescent live dyes.

Our results conclusively linked accumulating cytotoxic deoxySLs with mitochondria and endoplasmic reticulum (ER) damage triggering Autophagy of mitochondria and membrane cisterna of the ER. The induced autophagic flux ultimately leads to accumulating deoxySL containing intra-lysosomal lipid crystals. Lysosomal lipid substrate accumulation impaired physiological lysosome functions and caused cellular starvation. Lysosomal exocytosis appeared as a mechanism for cellular clearance of cytotoxic deoxySLs. In sum, our data define new ultra-structural targets of deoxySLs and link membrane damage to autophagy and abnormal lysosomal lipid accumulation. These insights may support new conclusions about diabetes type 2 and HSNA1 related tissue damage.

## INTRODUCTION

Lipids serve as essential functional components in cell membranes. The curvature of cell membrane can be dynamically changed by its lipid composition^1^. Dedicated lipid classes such as cholesterols or sphingolipids (SLs) are hallmarks for regulating lipid membrane microenvironments and shaping cell membrane curvature ^23^. An imbalance between different lipid species leads to aberrant membrane function and therefore to impaired cell metabolism and intracellular trafficking. Thus, lipid membrane alterations are frequently related to medical conditions such as mitochondrial disorders^4^, diabetes type 2 (T2DM) ^5^, neurodegeneration ^6^ or cancer^7^.

SL metabolism occurs in discrete subcellular membrane compartments and consist of a firmly regulated and interconnected metabolic network^8 9^. Cell viability is essentially connected to a dynamic balance between two antagonistic acting arms of SL metabolism: Intracellular accumulating ceramides (Cers) lead to cellular anti-survival effects (autophagy, apoptosis, growth inhibition), whereas an increased relative amount of sphinganine-1-phosphate (S1P) promotes cell survival (anti-apoptotic effects, cell motility, metastatic and drug resistant phenotypes). A firmly regulated metabolic conversion between both signaling arms of these bio-effector molecules is crucial to maintain cellular energy homeostasis and resulting tissue integrity ^10 11^. Therapeutic strategies for modulating ceramide levels in different subcellular compartments (plasma membrane, ER, mitochondria) for chemotherapies are already in use^12^. The implication of Cers in promoting insulin resistance, metabolic derangement and cell death was also reported ^13 14^. However, these general findings expose a whole network of questions concerning sub-cellular compartmentalization of SLs and related molecular mechanisms of different diseases. Cells contain numerous rare sphingolipid species potentially modulating relative sub-cellular sphingolipid levels^15^. One clinical relevant non-canonical class of SLs is 1-deoxysphingolipid (DeoxySL) that lacks the 1-hydroxygroup at the lipid head portion ^16^. DeoxySLs arise as a cellular product of SPTLC1 (serine palmitoyltransferase long chain base unit 1), due to a condensation reaction of alanine with palmitoyl-CoA to 1-deoxy-sphinganine (DOXSA). This deviates from the common condensation reaction of serine with palmitoyl-CoA which results in sphinganine (SA). Most enzymes, involved in SL and deoxySL *de novo* synthesis, are located to the ER ^17 18^. Like SA, DOXSA can be acetylated with fatty acids (FAs) to hydrophobic 1-deoxydihydroceramides (deoxyDHCers) by ceramide synthases^19 20^, followed by desaturation to 1-deoxy-ceramide (deoxyCer) ^21 22 23 24^. Due to the lack of the 1-hydroxgroup and a double bond in the head portion of deoxySLs, these lipids show increased hydrophobicity compared to its canonical counterparts ^21^. Moreover, limited lipid miscibility of deoxyCers resulting in separate gel like phases in GUVs (giant uni-lamellar vesicles) and putative forming of aggregates at higher concentrations was reported ^25^.

In cells the conversion of DeoxyCers to bioactive S1Ps or deoxySLs with complex head groups and export from the ER to the Golgi apparatus appear limited ^22^. Restricted intra-cellular transport and hindered catabolism of hydrophobic deoxySLs results in intracellular accumulation of deoxySLs in ER, mitochondria and lysosome associated compartments leading to organelle dysfunctions ^26 27 28 22^. Additionally, deoxySL-mediated disruption of the cytoskeleton was reported ^29^. Together these effects lead to susceptibility of energy demanding neurons for deoxySL-mediated ER-stress resulting in Unfolded Protein Response (UPR), mitochondrial degeneration and cytoskeletal changes in axons ^21 22 30^. Similar effects were also observed in proliferative cell lines, treated with DOXSA ^22^. A recent study reported tubular distorted mitochondria putatively containing deoxyCer/deox(DH)Cer after applying DOXSA on MEF cells. Additionally, by applying DOXSA increased autophagic flux and birefringent crystalline cellular lipid inclusions triggering NLRP3 inflammasome activation were also indicated ^31^. A clear identification of DOXSA-mediated autophagy-associated processes and fine-structural features of deoxySL-targeted organelles is still pending. DeoxySLs are of clinical relevance: Intra-cellular synthesis of deoxySLs is a hallmark for HSAN1 (hereditary sensory neuropathy type 1) and causes severe neurological problems in patients ^21^. In healthy human individuals a total blood plasm level of 0.1 to 0.3 μM DOXSA was indicated ^32 33 34^. In contrast, blood plasm levels of deoxySLs in HSAN1 can reach up to 1.2 μM ^21^; this value resonated with deoxySL levels in patients suffering from metabolic syndrome and diabetes type 2 ^33 34 35^. Complementary, elevated deoxySL levels were evident in streptozotocin-induced rat diabetes and leptin-deficient ob/ob mice ^36 33 37 38^. The toxic effect induced by intracellular accumulating deoxySLs plays also a role in cellular senescence in energy demanding cancer cells and deoxySLs were proposed as molecular intermediates in paclitaxel chemotherapy induced peripheral neuropathy ^29 28 39^. Accordingly, phase I clinical trials, investigating a potential anti-tumor effect of intravenously injected DOXSA, suggested deoxySL-mediated decrease of tumor size ^40 41 42 43^. However, severe neurodegenerative side effects related to DOXSA-therapy led to discontinuation of clinical trials.

Click-chemistry light microscopy (click-LM) lacks the resolution to determine all spatial aspects of lipid distribution on sub-organelle level and relies on antibody-based co-labeling approaches ^31 44 22^. Here, we complement the versatile tool box for detecting alkyne lipids ^45 22 46 47^ with a new labeling method for electron microscopy (EM). For realizing labeling of intracellular lipids in its complete ultra-structurally preserved biological context, we introduced a polyethylene glycol 1000 (PEG1000) functionalized 0.8 nm sized (ultra-small) gold-nanoparticle containing azide groups as reporter for sensitive one-step copper catalyzed alkyne-lipid detection. We termed this novel and detergent-free labeling approach “Golden-Click-Method” (GCM). We employed GCM for tracing alkyne-DOXSA ((2S,3R)-2-aminooctadec-17-yn-3-ol) and resulting metabolites in preserved sub-cellular microenvironments on the nanometer scale. Based on our ultra-structural studies, complemented with time-lapse light microscopical imaging, we propose that deoxySLs target the ER and mitochondria. Accumulating DeoxySL formed tubular lipid aggregates and compromised normal organelle structure and physiological functions. Accordingly, DeoxySL-associated mitophagy and ER-phagy of respective dysfunctional compartments was evident. Mitophagy and ER-phagy was accompanied by starvation induced non-selective autophagy. Different modes of DOXSA-mediated autophagy resulted in accumulating substrates including slowly degradable deoxySLs in expanded lysosomes. Observed intra-lysosomal buildup of lipid substrates, compromising lysosomal functions, was accompanied by lipid crystals storing deoxySLs. Lysosomal exocytosis, a process disposing accumulated lysosomal substrates, promoted cellular clearance after DOXSA-mediated lysosomal substrate accumulation. In sum, our findings revealed new facets of deoxySL-mediated damage of cell organelles, followed by mitophagy and ER-phagy, leading ultimately to detoxing cells via lysosome dependent pathways.

Categorizing targets of DOXSA and associated structural changes in a cell biological model system contributes to understand elementary sub-cellular processes and landmarks connected to DeoxySLs. Novel insights can contribute to reveal therapeutic strategies for counteracting deoxySL-mediated cytotoxicity, tissue damage and resulting neurodegenerative effects in relevant medical conditions such as HSAN1, chemotherapy induced neuropathy or diabetes type 2.

## METHODS

### Lipids / Alkyne-Lipids

Syntheses of alkyne-SA and alkyne-DOXSA are described^22 48^. DOXSA (1-deoxysphinganine) and Sphinganine (SA) were purchased from Avanti Polar Lipids (Alabaster, USA). Lipid were kindly provided by C. Thiele.

### General Cell Culture Procedures

Generating mouse embryonic fibroblast (MEF) cell line was described previously ^49^. MEF cells were maintained in DMEM medium (Sigma-Aldrich, Hamburg, Germany) containing 10 % fetal calf serum (FCS) (Sigma-Aldrich, Hamburg, Germany) and 1 % penicillin / streptomycin (Sigma-Aldrich, Hamburg, Germany).

### Detecting Apoptosis using Fluorescence Microscopy

For detecting apoptosis, MEF culture medium was supplemented with 1 μM DOXSA from ethanolic stock or vehicle (0.1 % ethanol). After incubating for 24 or 96 h, MEF cells were supplemented with CellEvent^™^ Caspase-3/7 green detection reagent (Life Technologies, Darmstadt, Germany) following manufacturer’s instructions before aldehyde fixation. DNA-counterstaining was done using DAPI-Fluoromount-G mounting medium (Science Services, Munich, Germany). Endpoint assays for detecting apoptosis were performed using confocal microscopy. Statistics were generated using Fiji ImageJ software^50^.

### Detecting Lipid Aggregates in MEF Cells using Fluorescence Microscopy

For detecting intra-cellular lipid aggregates, 70 % confluent MEF cell culture was supplemented with 1 μM DOXSA from ethanolic stock or vehicle (0.1 % ethanol). After incubating for 24 h, cells were treated with HCS LipidTOX Red Phospholipidosis detection Reagent (LipidTOX red) (Life Technologies, Darmstadt, Germany) according to manufacturer’s instructions. After aldehyde fixation cells were immediately imaged using LSM710 NLO in in µ-dish 35 mm imaging dishes with a glass cover slip bottom (Ibidi, Munich, Germany).

### Analysis of Subcellular Localization of Alkyne-Lipids in MEF cells using Fluorescence Microscopy

For fluorescent detection of intra-cellular alkyne-deoxySLs, cells were grown on glass cover slips until 70 % percent confluence was reached. Then MEF cells were supplemented with 1µm alkyne-SA or 100 nM alkyne-DOXSA and 900 nM DOXSA from ethanolic stocks. After 24 h cells were fixed and fluorescent click- and antibody labeling was performed as described previously^22^.

Imaging was performed using an LSM710 confocal laser-scanning microscope equipped with a Transmitted Photomultiplier Tube (T-PMT) and controlled by the software ZEN 2012 (Carl Zeiss, Jena,= Germany) was used for confocal fluorescence and bright field microscopy. Samples were imaged using an EC Plan-Apochromat 63x/1.4 oil DIC M27 objective. Images were deconvolved with Huygens Professional version 19.04 (Scientific Volume Imaging, The Netherlands, http://svi.nl).

### Analysis of Membrane Dynamics in MEF cells using Fluorescent Time-Lapse Imaging

For time-lapse (TL) imaging of organelle dynamics, MEF cells were cultured in µ-dish 35 mm imaging dishes with a glass cover slip bottom (Ibidi, Munich, Germany) until 70 % confluence. Cell culture medium was supplemented with organelle markers (ER-Tracker Blue-White [Ex.: 374 nm / Em. 430 - 640 nm nm], MitoTracker Green FM [Ex.: 490 nm / Em. 516 nm] or LysoTracker Red DND-99 [Ex.: 577 nm / Em. 590 nm]) (Life Technologies, Darmstadt, Germany) according to manufacturer’s instructions. Before TL-imaging, cells were supplemented with 1 µM DOXSA from ethanolic stock solutions or vehicle (0.1 % ethanol).

The glass bottom dish was mounted on a spinning disc (SD) confocal microscope setup (Cell Observer SD Spinning Disk, controlled by the software ZEN 2012 [Carl Zeiss, Jena, Germany]) and kept at a steady temperature of 37 °C with 5 % CO_2_. TL-imaging was performed using a Plan-Apochromat 63 × / 1.4 oil objective. Statistics were generated using Fiji ImageJ software^50^.

### Preparing Adherent Cells for Scanning Electron Microscopy

Cells grown on 15 mm glass cover slips were supplemented with 1 µM DOXSA from ethanolic stock or vehicle (0.1 % ethanol) for 24 h, before fixing with 2 µM Paclitaxel, 4 % PFA and 2.5 % GA in 50 mM cacodylate buffer (Sigma-Aldrich, Hamburg, Germany) pH 7.2 (CB) for 10 min at 37 °C. Critical point drying (CPD) was performed using a CPD 030 (BAL-TEC, Balzers, Liechtenstein) after dehydration with ethanol. Cover slips were mounted on aluminum sample holders using conductive silver paint (Plano, Wetzlar, Germany) before sputter coating with 2 nm platinum/palladium in an HR 208 coating device (Cressington, Watford, UK) and imaging with an FEI Verios 460L (FEI, Eindhoven, The Netherlands) using a through-lens low energy electron detector (TLD) at 5 kV acceleration voltage.

### Analysis of Ultra-Structural Localization of Alkyne-Lipids in MEF cells using Golden Click Method (GCM)

For alkyne-lipid labeling using GCM MEF cells supplemented with 100 nM alkyne-DOXSA and 900 nM DOXSA for 24 h were fixed using 4 % PFA and 2.5 % GA in 50 mM HEPES for 20 min at 37 °C (pH 7.45) (for morphological examinations 50 mM CB [pH 7.2] was used). Subsequently cells were rinsed with fresh fixative and buffer.

PEG-1000-azide ultra-small particles (custom conjugation by Aurion, Wageningen, The Netherlands) were diluted in 50 mM HEPES (+ 150 mM NaCl) to a final concentration of 5 × 10^12^ particles/ml; to 4 °C pre-cooled gold particle solution was transferred to samples. The click-reaction was initiated by adding of 2.3 mM Cu(I)TFB in acetonitrile (final 2.3 % acetonitrile) and performed at 4 °C for 48 h. After alkyne-lipid labeling, samples were extensively washed using 50 mM HEPES (+ 150 mM NaCl) at room temperature (RT). For visualizing ultra-small gold reporters in the EM the Silver Enhancement for Electron Microscopy Kit (SE-EM) (Aurion, Wageningen, The Netherlands) was applied for 35 min according to the manufacturer’s instructions. Before SE-EM, cells were transferred to Enhancement Conditioning Solution (ECS) (Aurion, Wageningen, The Netherlands). After SE-EM, cells were thoroughly rinsed using ECS and transferred to CB before further sample processing for TEM.

Post-fixation of lipids with 0.1 % Malachite Green and 2.5 % glutaraldehyde in CB was done for 16 h at 4 °C. Afterwards cells were twice washed using CB containing 200 mM sucrose and twice with CB. Next, cells were contrasted by using 1 % osmium tetroxide (OsO_4_) (Science Services, Munich, Germany) and 0.8 % potassium hexacyanoferrate (III) (Sigma-Aldrich, Hamburg, Germany) in CB for 1 h on ice followed by rinsing with CB on ice. A subsequent second fixation using 1 % OsO_4_ in CB was done for 1 h on ice. After washing with CB and water, the samples were treated with 1 % tannic acid (TA) in water for 10 min at RT. After rinsing thoroughly with pure water, cells were further contrasted using 1 % uranium acetate (UA) in water for 1 h at 4 °C. Sample dehydration using an ascending ethanol row; 20 min final dehydration in 100 % water free ethanol; embedding using Epon substitute embedding kit (Sigma-Aldrich, Hamburg, Germany) and polymerizing for 2 -3 d at 60 °C followed.

Whole dishes were mounted in the LSM 710 NLO for visualizing the silver enhanced GCM labeling using laser reflection microscopy in a broader perspective. By silver grains reflected light was detected with the smallest possible setting for pinhole diameter in “reflection mode”.

The glass bottom of the culture dish was removed by dipping the dish into liquid nitrogen before pre-trimming and mounting onto a Leica UC7 (Leica Microsystems, Wetzlar, Germany). Before generating 65 nm ultra-thin sections using an ultra 35° diamond knife (Diatome AG, Biel, Switzerland), a trapeze shaped pyramid (400 × 200 µm) was trimmed using a Trim 90 trimming diamond knife (Diatome AG, Biel, Switzerland). Longitudinal plastic sections of adherent MEF cells were mounted on 1 × 2 mm slot grids with carbon coated formvar film (Plano, Wetzlar, Germany).

Ultra-thin sections were analyzed using scanning transmission electron microscopy (STEM) and electron tomography (ET). STEM was performed with a FEI Verios 460L using a transmission high energy electron detector at 25 kV acceleration voltage. Tomograms for high resolution analysis were acquired with a JEOL JEM 2200 FS TEM with an omega energy filter and an acceleration voltage of 200 kV (JEOL, Japan) equipped with a Temcam-F416 (TVIPS, Munich, Germany). The omega energy filter slit was set to 20 eV. Tilt-acquisitions (from -70° to + 70°) were obtained using SerialEM software ^51^. Tomograms were reconstructed in eTomo, part of the IMOD software package^52^ using patch tracking mode and weighted back projection (WBP). Statistics were generated using Fiji ImageJ software^50^.

## RESULTS

### The Golden Click Method (GCM): A New Method for Detecting Alkyne-Lipids on Nanometer Scale Using Azide-Gold-Reporters

Intracellular lipid trafficking and firmly regulated lipid compartmentalization are indispensable for cellular physiology^53^. Perturbating cellular lipid transport and homeostasis on sub-cellular level can lead to cellular malfunctions resulting in disease. Thus, accurate spatial and metabolic tracing of intra-cellular lipids on ultra-structural level is of emerging interest to the scientific community.

Here, we present a novel one-step labeling approach for detecting virtually native acting alkyne-lipid analogues on the nanometer scale. The triple-bond (alkyne-group) is one of the smallest molecular tags; it virtually does not alter the biophysical properties of the lipid, while enabling sensitive tracing via a copper(I)-catalyzed azide alkyne cycloaddition (“click-reaction”) (Fig. 1 A). Previous studies introduced robust protocols that allow the use of alkyne lipids to quantify lipid metabolism biochemically, using mass spectrometry and to trace them spatially applying fluorescence microscopy ^45 44 46 54 31^. Our work complements the established toolbox for detecting alkyne-lipids with a small amphiphilic gold-containing probe readily diffusing into aldehyde fixed cells, suitable for sensitive lipid tracing on ultra-structural level (Fig. 1 B). We termed this new labeling approach “Golden-Click-Method” (GCM). After click-labeling of alkyne-deoxySLs with functionalized ultra-small gold-nanoparticles, a silver-enhancement for EM was performed before sample processing. Electron-dense silver particles (8 - 10 nm) were detected in cells treated with alkyne tracer but were largely missing in control samples (Fig. 1 C - C’’). Specific alkyne-deoxySL labeling with metallic reporters was confirmed in a broader perspective using confocal reflection microscopy (FIG. 1 D + D’).

**Figure 1:**
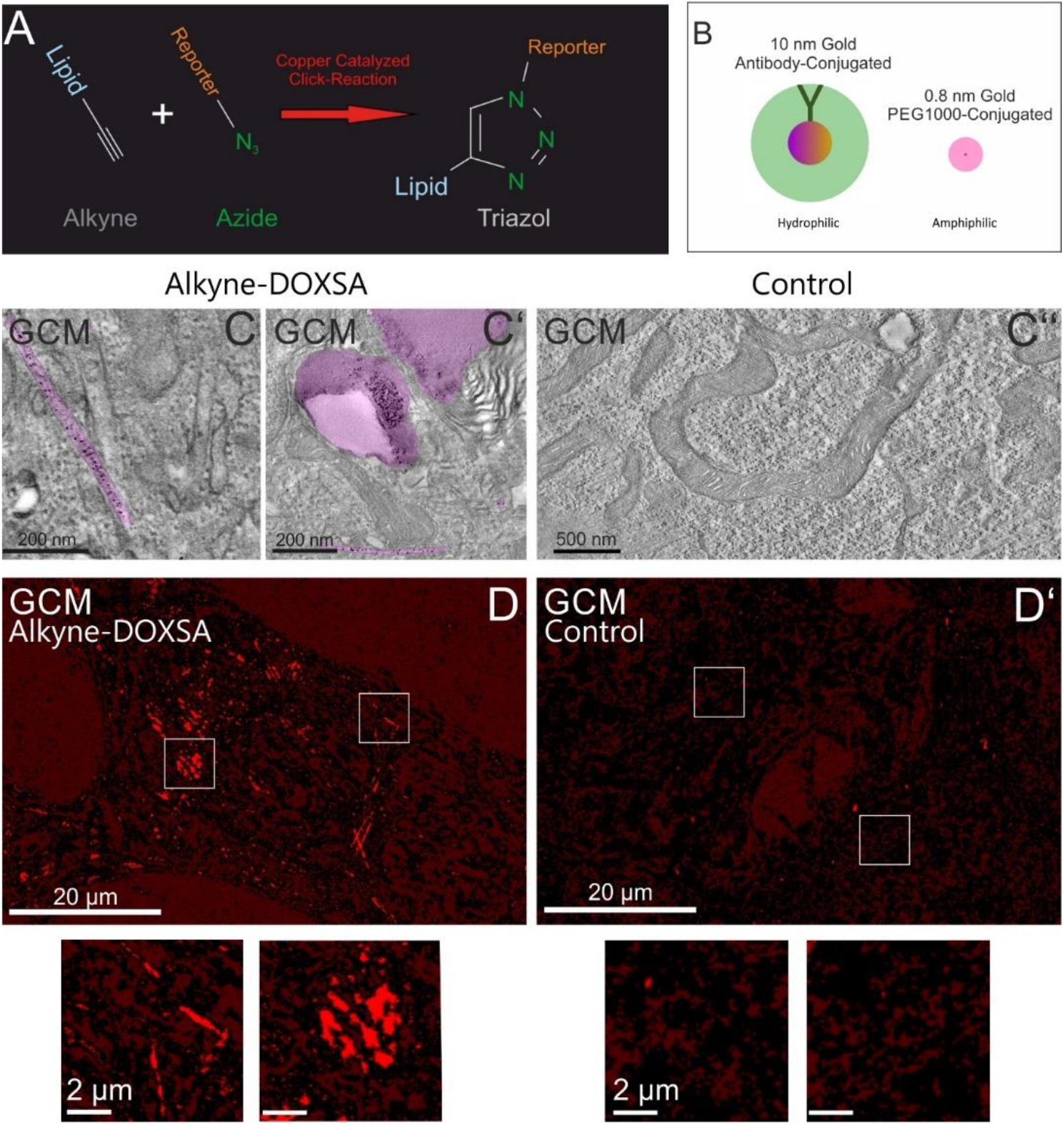
The Golden Click Method (GCM): A new method for detecting alkyne-lipids on nanometer scale using azide-gold-reporters. A) Scheme representing the chemical reaction for copper(I)-catalyzed azide alkyne cycloaddition (“click-reaction”) for detecting alkyne-lipids. B) Amphiphilic azide-PEG1000-functionalized ultra-small gold-probe (right) compared to a hydrophilic antibody-conjugated 10 nm gold particle (left). C) MEF cells were supplemented with 100 nM traceable alkyne-DOXSA and 900 nM DOXSA (or vehicle) for 24 h followed by chemical fixation, pre-embedding labeling using Golden Click Method (GCM) and silver enhancement for EM. While alkyne-containing cells comprised electron dense labeling in discrete intra-cellular compartments (magenta), vehicle treated cells exhibited low unspecific background labeling. D) Metallic labeling in plastic embedded cells was verified in a broader perspective using confocal reflection microscopy. Inserts (white boxes) show representative signals in detail (LUT: Black-to-Red, Zeiss ZEN).

### Accumulating DeoxySLs Interfere with Cell Proliferation

Previously, DeoxySLs were shown to interfere with lipid membranes, mitochondrial health and cell proliferation ^31 22 25^. Using bright field microscopy, we confirmed changes in internal cell morphology and cell polarity after exogenously applying 1 µM DOXSA for 24 h to mouse embryonic fibroblasts (MEF cells) (Fig. 2 A + A’). Additionally, DNA labeling using DAPI (4′,6-Diamidine-2′-phenylindole dihydrochloride) of equal treated cells showed an increase in multinucleated cells from 2.3 to 12.7 % corroborating by DOXSA impaired cell division (n_24h DOXSA_ = 1467; n_24h Control_ =2382) (Fig. 2 B). Consequently, DOXSA-treated MEF cells exhibited by approximately 50 % reduced growth density compared to control cells after 24 h. 96 h after applying DOXSA, growth density appeared recovered indicating a transient anti-proliferative effect of DOXSA on MEF cells (Fig. 2 C). For excluding deoxy-SL mediated apoptosis, we applied CellEvent^™^ Caspase-3/7 Green detection reagent (CellEvent) together with DAPI to equal treated cells. We performed an endpoint assay using LM revealing a minor increase in apoptosis from < 0.1 to 3 % after applying 1 µM DOXSA (n_24h DOXSA_ = 1467; n_24h Control_ =2382; n_96h DOXSA_ = 2290) for 24 h and 96 h (Fig. 2 D). Thus, we suggest a minor role of apoptosis in deoxySL associated tissue damage.

**Figure 2:**
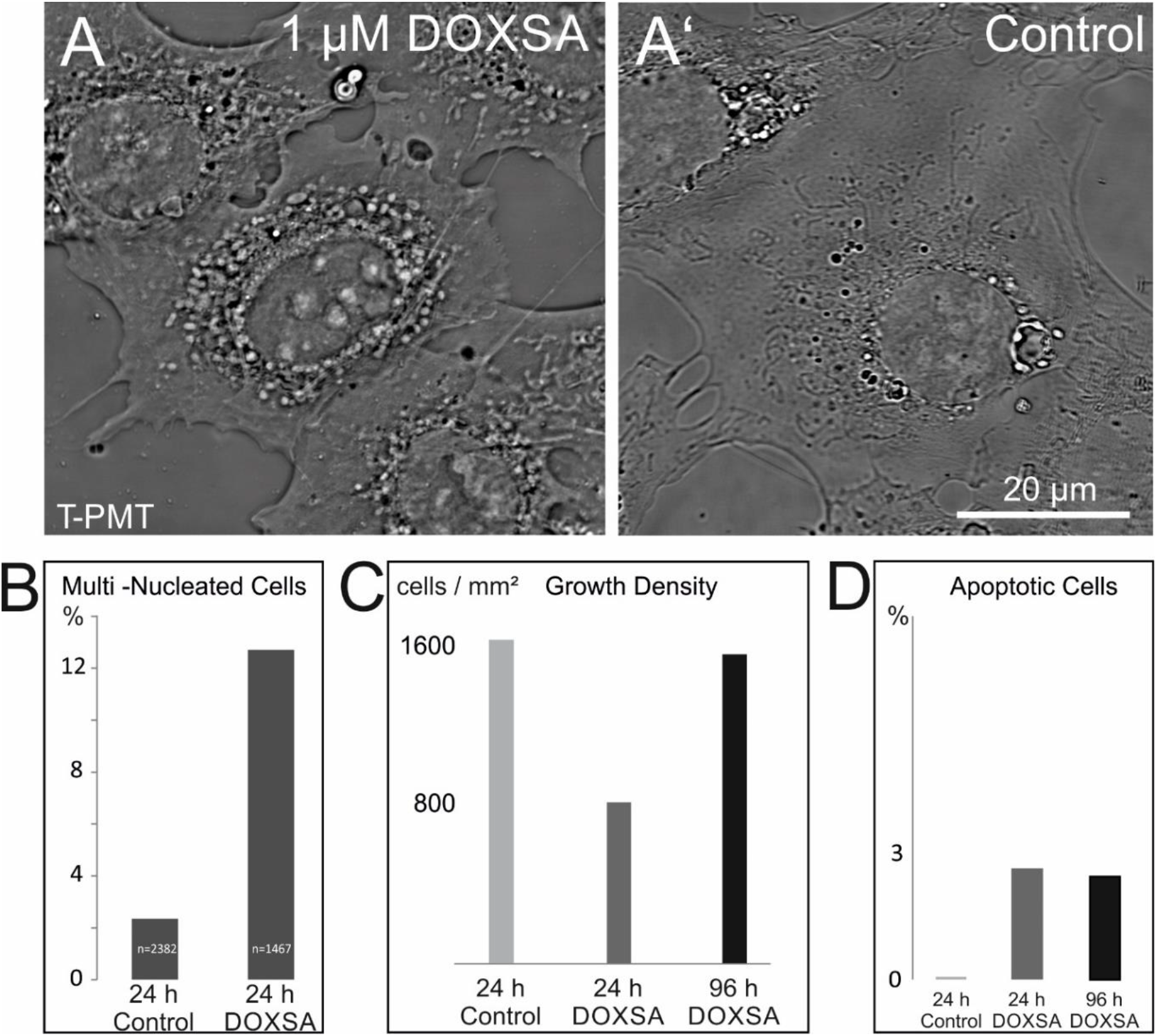
Accumulating deoxySLs interfere with cell proliferation. A+A’) MEF cells were supplemented with 1 µM DOXSA or vehicle (0.1 % EtOH); bright field microscopy (T-PMT) indicated sub-cellular membrane alterations and impaired cell polarity after applying DOXSA. B) Cells were equal treated before DAPI staining. The endpoint assay indicated a 6-fold increase of multi-nucleated cells compared to control. C) 24 h after applying DOXSA, density of MEF cells appeared by 50 % decreased compared to control; 96 h after application growth density appeared recovered. E) CellEvent reagent revealed relatively low levels of apoptosis after applying DOXSA to MEF cells.

To investigate the effect of deoxySLs on sub-cellular membrane compartments we performed EM on longitudinal sections of DOXSA-treated MEF cells. MEF cells revealed extended tubular lipid-rich compartments (arrowheads) associated with electron-opaque lipid aggregates (size: 0.5 – 2 µm), fragmented mitochondria showing cristae damage and abundant auto-lysosomal compartments with putative high lipid content. Nuclei (N) and cell shape appeared often disfigured (Fig. 3 A + B, S1 A + A’, S2 A-A’’). An endpoint assay using LipidTox Red Phospholipidosis Detection Reagent (LipidTox) verified high lipid content in extended (arrowheads) and spherical compartments (Fig. 3 C +D).

**Figure 3:**
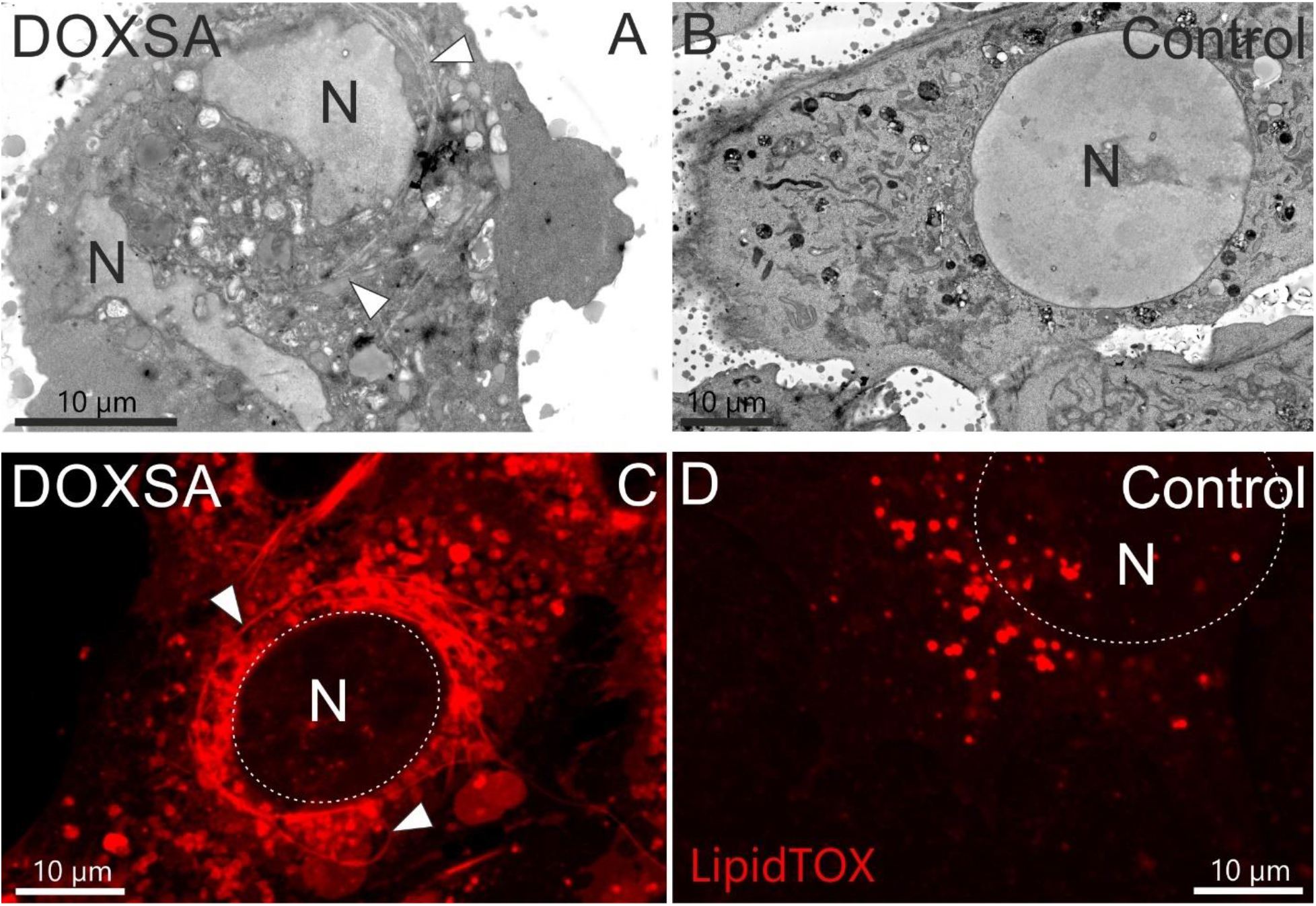
DeoxySLs tempted lipid accumulation in MEF cells. A+B) Ultra-structural examination of MEF cells exhibited cell compartments associated with lipid-rich inclusions after applying 1 µM DOXSA. Nuclei (N) and cell shape appeared often disfigured. C + D) High lipid content of extended and spherical inclusions (arrowheads) was corroborated using LipidTox Phospholipidosis Detection Reagent (dashed line indicates nucleus).

Completive, we re-investigated cells treated with 900 nm DOXSA + 100 nm alkyne-DOXSA for 24 h using deconvoluted confocal microscopy. In accordance to previous studies ^22 31^, alkyne-DOXSA targeted mitochondria, spherical structures and different extended tubular compartments. Different sub-cellular alkyne-lipid labeling patterns in adjacent cells indicated putative cell-cycle depended differences in DOXSA-metabolism resulting in different stages of DOXSA-intoxication. MEF cells treated with 1 µM Sphinganine (SA) (900 nM SA + 100 nm alkyne-SA, 24 h) indicated alkyne-SL labeling mostly at mitochondria (Fig. S3 +S4).

### DeoxySLs Aggregated in Smooth ER and Mitochondria

Most enzymes, involved in SL and deoxySL *de novo* synthesis, are located to the ER ^17 18^. This results in aggregating hydrophobic deoxyCers in the ER-compartment, where these lipids are putatively generated from its precursor DOXSA. To understand the effect of deoxySLs on ER and mitochondrial membranes, we traced alkyne-deoxySLs in its ultra-structurally preserved microenvironments using GCM.

In accordance to click-LM (Fig. S3), we recognized alkyne-positive tubular assemblies in the perinuclear area using EM. Based on absence of ribosomes and the rough ER (rER) marker PDI from tubular compartments, we suggest aggregating deoxySLs in the smooth ER compartment (sER). GCM labeling for alkyne-deoxySLs revealed a clear-cut localization of alkyne-deoxySL to the electron opaque lumen of indicated tubes (red, arrowheads) and associated lipid particles (magenta, asterisk) (Fig. 4 A + B). Examining further EM-samples using ET defined the fine structural characteristics of lipid-rich tubes (arrowhead), which appeared continuous with the rER (yellow). We observed an electron opaque lipid-rich core (diameter 45 - 50 nm) bordered by a lipid membrane monolayer (Fig. 4 C + C’, Fig. 5 E).

**Figure 4:**
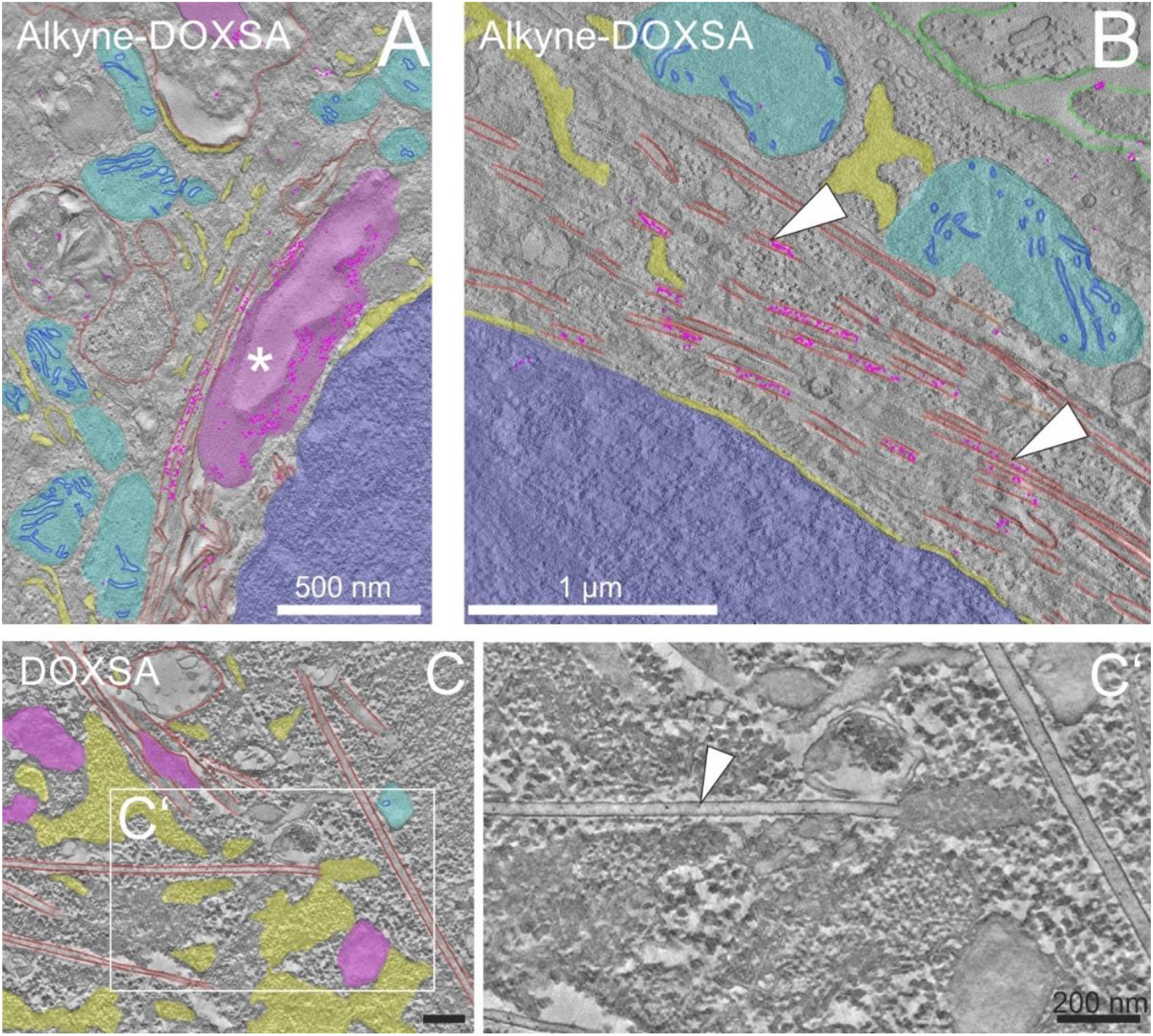
DeoxySLs aggregated in smooth ER. A) Electron tomography (ET) of GCM confirmed alkyne-deoxySL labeling (pink dots) in lipid aggregates (magenta /asterisk) and B) in putative smooth ER-derived tubes (red / arrowheads) after applying 900 nM DOXSA and 100 nM alkyne-DOXSA. C) ET of MEF-cells, treated with 1 µM DOXSA, indicated connections between tubes (red) and ER-domains (yellow). C’) Zoom-in from C) without overlay, ER-derived tube is labeled with an arrowhead. (ER-derived tube = red; lipid aggregate = magenta, nucleus = blue; mitochondria = cyan, plasma membrane = green, GCM labeling = pink dots).

**Figure 5:**
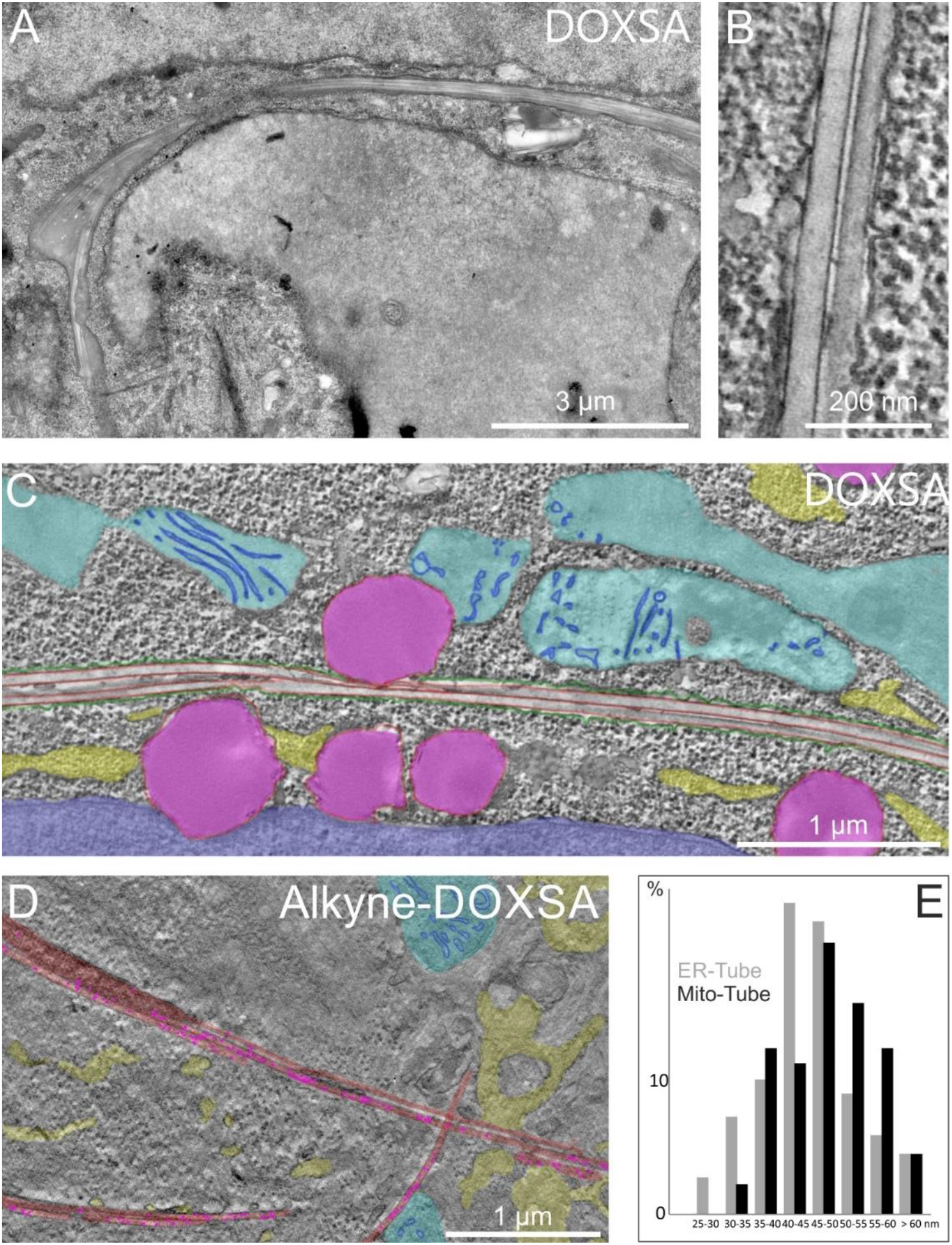
DeoxySLs aggregated in mitochondria. A) After applying 1 µm DOXSA, EM indicated extended mitochondria including tubular lipid inclusions. Contact with lysosomes was evident. B) ET revealed the fine structure of tubular inclusions. C) Tubular mitochondria were frequently associated to small lysosomes (magenta) D) After applying 900 nM DOXSA and 100 nM alkyne-DOOXSA, GCM confirmed a clear-cut localization of alkyne-deoxySL labeling (magenta dots) to the lumen of intra-mitochondrial tubes (red). E) Quantitative EM indicated similar characteristic tube diameters at ER and mitochondria. (mitochondria = cyan; ER = yellow; intra-mitochondrial tubes = red; lipid particles = magenta; nucleus = blue; outer mitochondrial membrane of tubular mitochondrium = green; GCM labeling = pink dots)

Import of deoxySLs into mitochondria was previously reported ^22 31^. Accordingly, our EM-data exhibited elongated mitochondria comprising bundles of tubular lipid-inclusions apparently connected with mitochondrial cristae. Interplay between lysosomes and tubular mitochondria was evident (Fig. 5 A). Mitochondrial inclusions (red) appeared with an average diameter of 45 - 50 nm like ER-related lipid tubes (Fig. 5 B +C, Fig. 5 E) and were also positive for GCM labeling of alkyne-deoxySLs (Fig. 5 D). Thus, we suggest similar deoxySL-induced membrane disturbing effects in ER and mitochondria.

To exclude biological artifacts induced by applying lipids to MEF cells, the fine-structure of DOXSA-treated cells was compared to vehicle controls treated with 0.1 % ethanol. To exclude structural changes induced by a high lipid load, MEF cells, treated with 1 µM sphinganine (SA) or 20 µM oleate (Ole), were examined. In sum, our controls appeared inconspicuous. Details can be found in figures S1 and S2.

### DeoxySLs Tempt Mitophagy

Previous studies indicated that exposure of MEF cells and neurons to DOXSA tempted cellular stress leading to mitochondrial fission and autophagy ^22 31^. However, temporal and ultra-structural aspects of interplay between autophagic processes targeting mitochondria and deoxySLs were not assessed. We re-investigated DOXSA-induced mitochondrial fission using time-lapse (TL) imaging with MitoTracker Green FM (MitoTracker) (Video S1). Quantitative TL-imaging specified an almost linear rate of DOXSA-mediated mitochondrial fission over time until the mitochondrial network appeared entirely fragmented after about 4.5 h (Fig. 6 A + B). Additionally, live cell imaging revealed individual mitochondrial fission events (pink asterisk) as a rapid process faster than 1 second (Fig. 6 C, Video S2). EM on corresponding MEF cells confirmed DOXSA-mediated mitochondrial fission resulting in separated mitochondrial compartments (yellow asterisk). A portion of structurally disfigured mitochondria was interacting with auto-lysosomal compartments (red asterisk) (Fig. 6 D + D’). Tracing alkyne-deoxySLs in equal treated cells on ultra-structural level confirmed interplay of deoxySLs with mitophagy. GCM indicated a high deoxySLs in structurally disfigured mitochondria interacting with auto-lysosomal compartments (Fig. 6 E-F). Mitochondrial fission together with mitophagy may display a protective mechanism separating deoxySL-intoxicated dysfunctional from functional mitochondrial sub-compartments supporting cellular regeneration after applying DOXSA. Accordingly, GCM labeling was rarely detected in structurally intact mitochondrial cross-sections.

**Figure 6:**
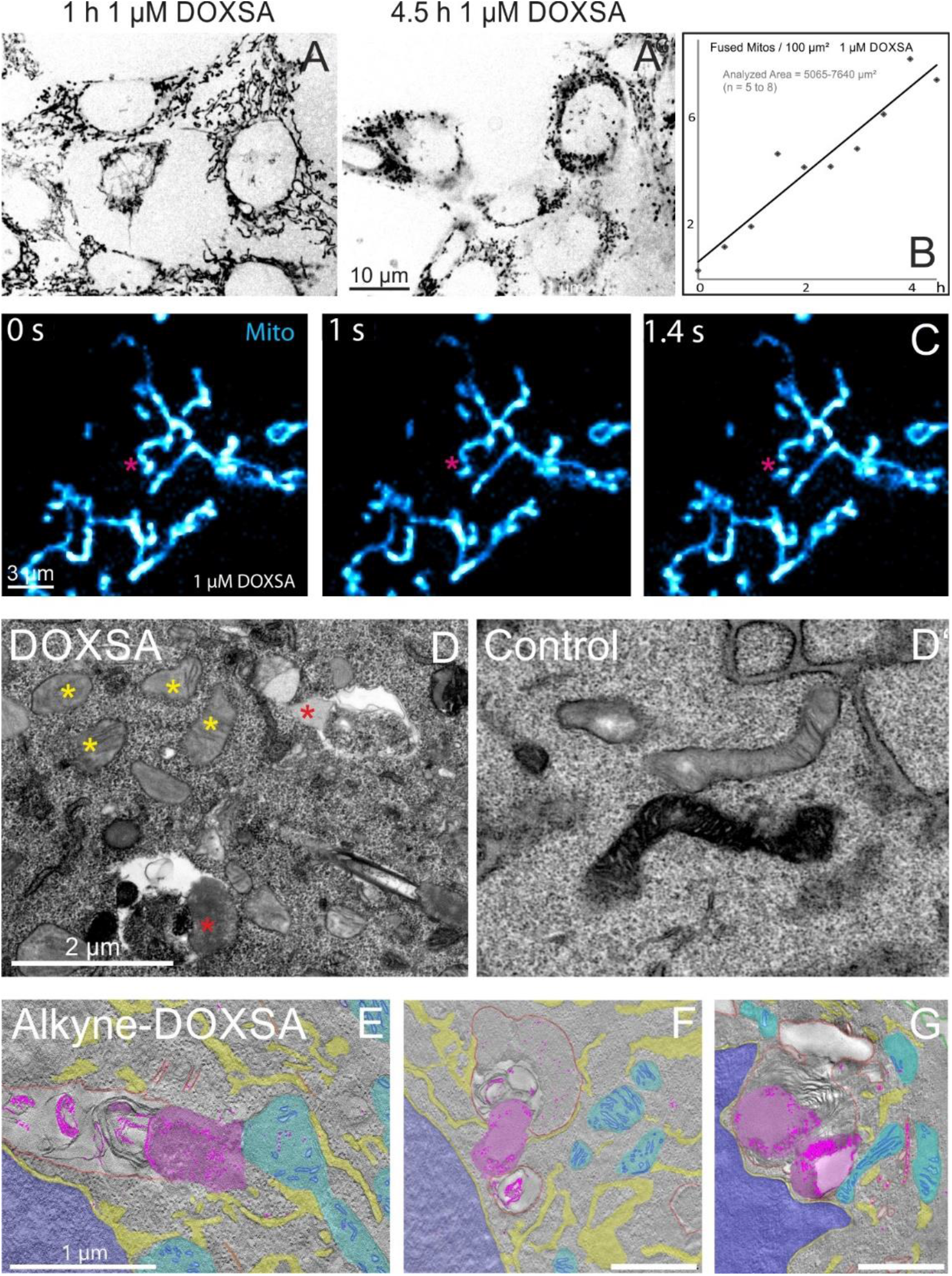
DeoxySLs tempt mitophagy. A+A’) Mitochondrial fission was observed after applying 1 µM DOXSA to MEF cells. B) TL-imaging with MitoTracker confirmed disintegration of the mitochondrial network over time. C) Individual DOXSA-mediated mitochondrial fission events (pink asterisk) were tracked using spinning disc microscopy of MitoTracker (5 fps). D + D’) EM showed mitochondrial fission resulting in compartments including intact cristae (yellow asterisk) and putative dysfunctional subunits interacting with autolysosomal compartments (red asterisk). E-G) After applying 900 nM DOXSA and 100 nM alkyne-DOXSA, GCM confirmed interplay between impaired mitochondria (cyan), autolysosomal compartments (red) and alkyne-deoxySLs (magenta dots). Mitophagy was associated with alkyne-positive mitochondrial remnants (magenta) (ER = yellow; nucleus = blue; GCM labeling = pink dots)

### DeoxySLs Tempt ER-Phagy and Starvation Induced Autophagy

Elevated levels of deoxySLs were shown to cause ER-swelling, ER-collapse and ER-stress ^22 31^. Indicated effects point on ER-phagy involved in the cellular response on DOXSA-intoxication. However, details about deoxySL-mediated ER-phagy remained largely unknown.

To follow the fate of ER-membranes after applying 1 µM DOXSA to MEF cells, we performed TL-imaging of the fluorescent ER-tracker blue/white DPX (ER-tracker) (Video S3). In addition to ER-collapse, we recognized shaping of hollow-like ER-tracker positive compartments after about 3 h in the perinuclear area (green arrowhead). After approximately 5 – 6 h ER-tracker positive spherical compartments (pink arrowhead) appeared close to straight elongated compartments (Fig 7 A’, A’’ + B).

**Figure 7:**
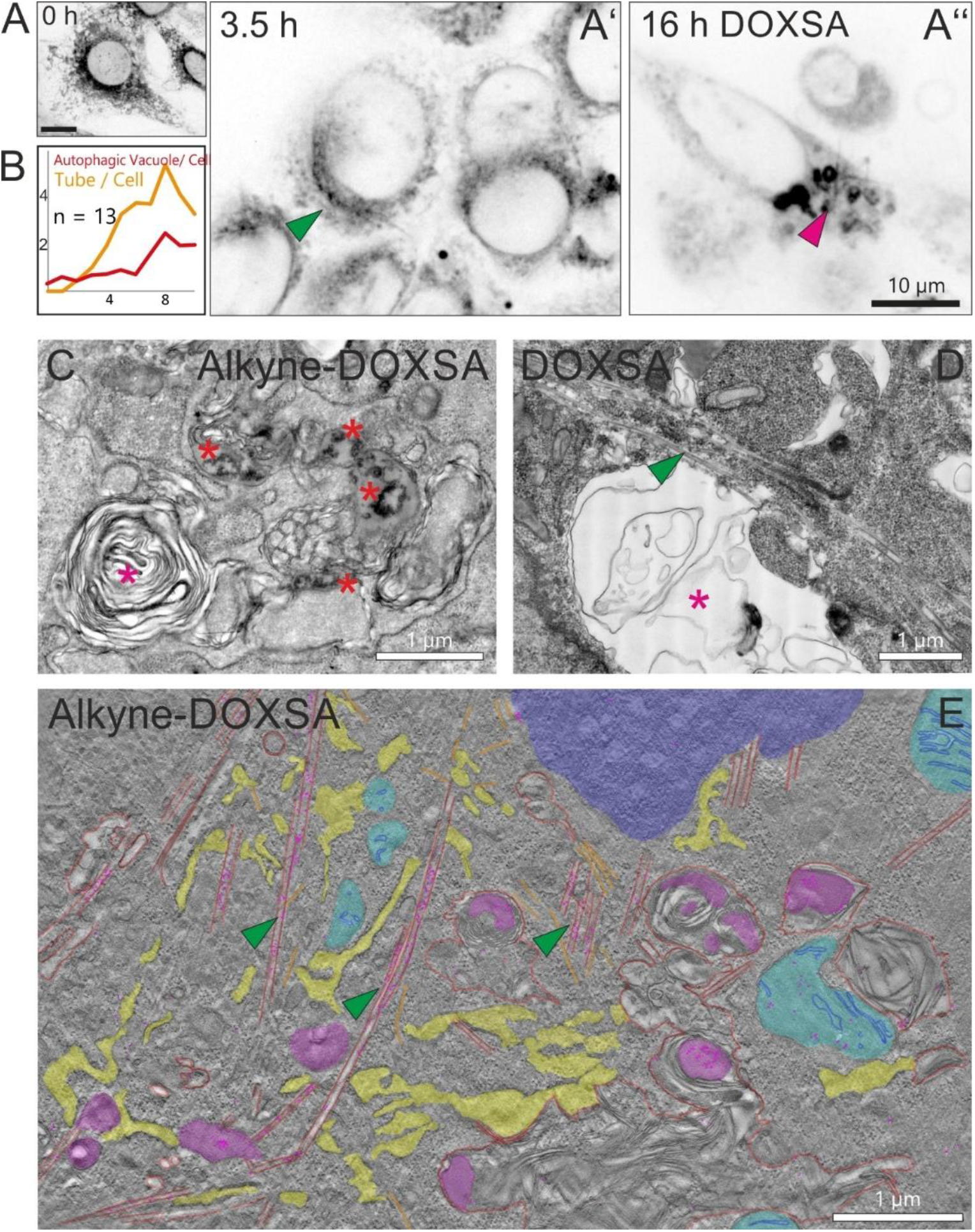
DeoxySLs tempt ER-phagy. A) ER Tracker Blue-White DPX (ER-tracker) was applied to MEF cells before applying 1 µM DOXSA. A’) Resulting TL-imaging demonstrated forming of hollow appearing assemblies positive for ER-tracker (green arrowhead) after DOXSA-induced ER-collapse. A’’) The fluorescent ER-tracker was frequently translocated to spherical compartments and straight structures (pink arrowhead). B) Quantifying TL-acquisitions confirmed forming of hollow like assemblies after 2-3 h; an increase in spherical compartments together with stiff tubes was indicated after 6-7 h. C) Examining MEF cells, treated with 900 nM DOXSA and 100 nM alkyne-DOXSA, revealed ultra-structural characteristics of ER-phagy including membrane containing vesicles (magenta asterisk) associated with alkyne-deoxySL labeling (red asterisk). D) Additionally, interaction of membrane containing vesicles (magenta asterisk) with ER-derived tubes (green arrowhead) was observed after applying 1 µM DOXSA and E) applying 900 nM DOXSA and 100 nM alkyne-DOXSA. (ER = yellow, ER-derived tube and related autolysosome = red; lipid aggregate = magenta, nucleus = blue; mitochondria = cyan; GCM labeling = pink dots).

Examining corresponding MEF cells using GCM revealed proximity between disfigured ER-domains associated to alkyne-positive lipid aggregates (red asterisks) and membrane containing vesicles (pink asterisk) corroborating DOXSA-mediated ER-phagy (Fig 7 C). Additionally, EM revealed membrane containing autolysosomes (red asterisk) in close contact with ER-derived deoxySL-rich lipid-tubes (green arrowhead) (Fig. 7 D). GCM confirmed alkyne-deoxySL labeling at ER-derived tubes (red, green arrowheads) interacting with lysosomal compartments (red) (Fig. 7 E).

Next, we aimed to investigate straight tubes positive for the ER-tracker which appeared 5 - 6 h after applying DOXSA. EM revealed membrane tubes with a diameter of 200 – 250 nm bordered by 3 - 6 stacked lipid membranes (pink arrowhead) connected to autolysosomal compartments (red asterisk) (Fig 8 A). Indicated membrane tubes (red) contained alkyne-positive lipid particles (magenta) together with cytoplasmic constituents pointing on its involvement in starvation induced autophagy (Fig. 8 B + C) ^55 56^. A cross-section of an incomplete formed membrane tube (pink arrowhead) showed ultra-structural characteristics of an isolation-membrane engulfing mitochondria (asterisk) together with surrounding cytoplasm (Fig 8 D).

**Figure 8:**
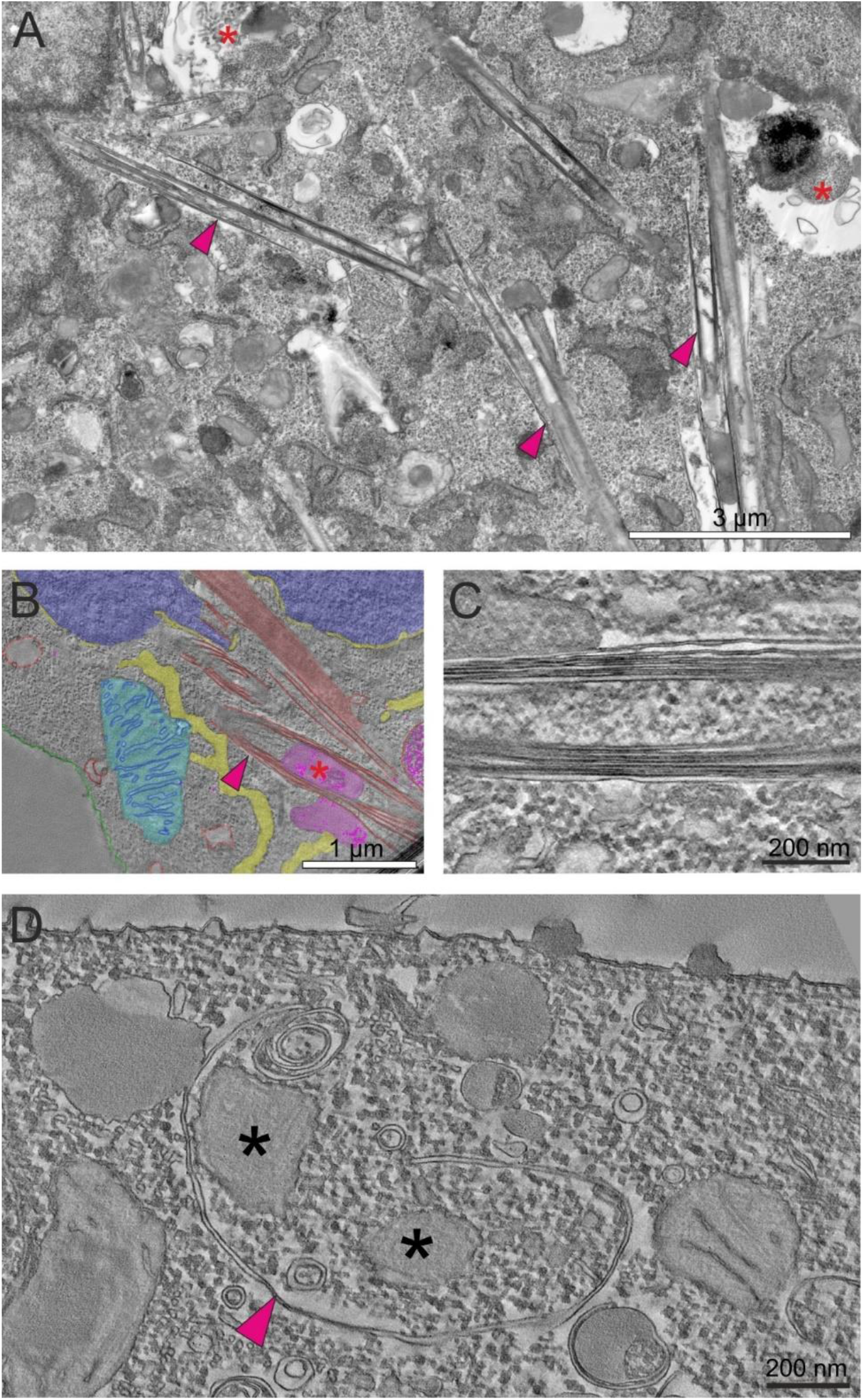
DeoxySLs tempt starvation induced autophagy: A) After applying 1 µM DOXSA, MEF cells revealed straight tubular compartments (pink arrowhead) closely associated to autolysosomal compartments (red asterisk). B) After treatment with 900 nM DOXSA and 100 nm alkyne-DOXSA, GCM indicated alkyne-deoxySLs in lipid particles (magenta) enclosed together with cytoplasmic constituents in membrane tubes (red). C) ET confirmed cytoplasm enclosed into 3 – 6 stacked lipid membranes. C) A cross-section of an incomplete formed membrane tube corroborated ultra-structural characteristics of an isolation membrane engulfing mitochondria and cytoplasm. (nucleus = blue; mitochondria = cyan; ER = yellow; GCM labeling = pink dots)

### DeoxySLs Accumulate in Lysosomes

Previously birefringent lipid crystals, presumably containing deoxyCers and deoxy(DH)Cers, were reported after applying DOXSA to MEF cells ^31^. However, dynamics, fine-structure and spatial relation of lipid crystals to the auto-lysosomal system remained largely undescribed. Therefore, we investigated the effect of applying 1 µm DOXSA on the lysosomal content in MEF cells. Examining DOXSA-treated MEF cells using EM indicated expanded lysosomal compartments (pink asterisk) comprising lipid particles, lipid membranes and cytoplasmic debris while lysosomes in control sections (pink asterisk) appeared small and unremarkable (Fig. 9 A + B). Quantitative EM indicated increased contact of mitochondria with lysosomes and corroborated deoxySL-induced lysosomal expansion (Control: 2.6 % of diameters > 1500 nm; 1 µM DOXSA 24 h: 42.4 % of diameters > 1500 nm) (Fig. 9 C + D). We re-investigated crystalline lysosomal inclusions on fixed MEF cells using a co-staining of MitoTracker (cyan) with lipophilic LipidTox dye (red). Needle-shaped lipid containing inclusions reached up to 15 µm in length and appeared isolated or in bundles (Fig 9 E). EM revealed corresponding lipid crystals (green) in expanded lysosomes (rose) of equal treated MEF cells. Elongated lipid crystals were surrounded by lipid membrane bulks, lipid particles (magenta) and cytoplasmic remnants (Fig. 9 F + G).

**Figure 9:**
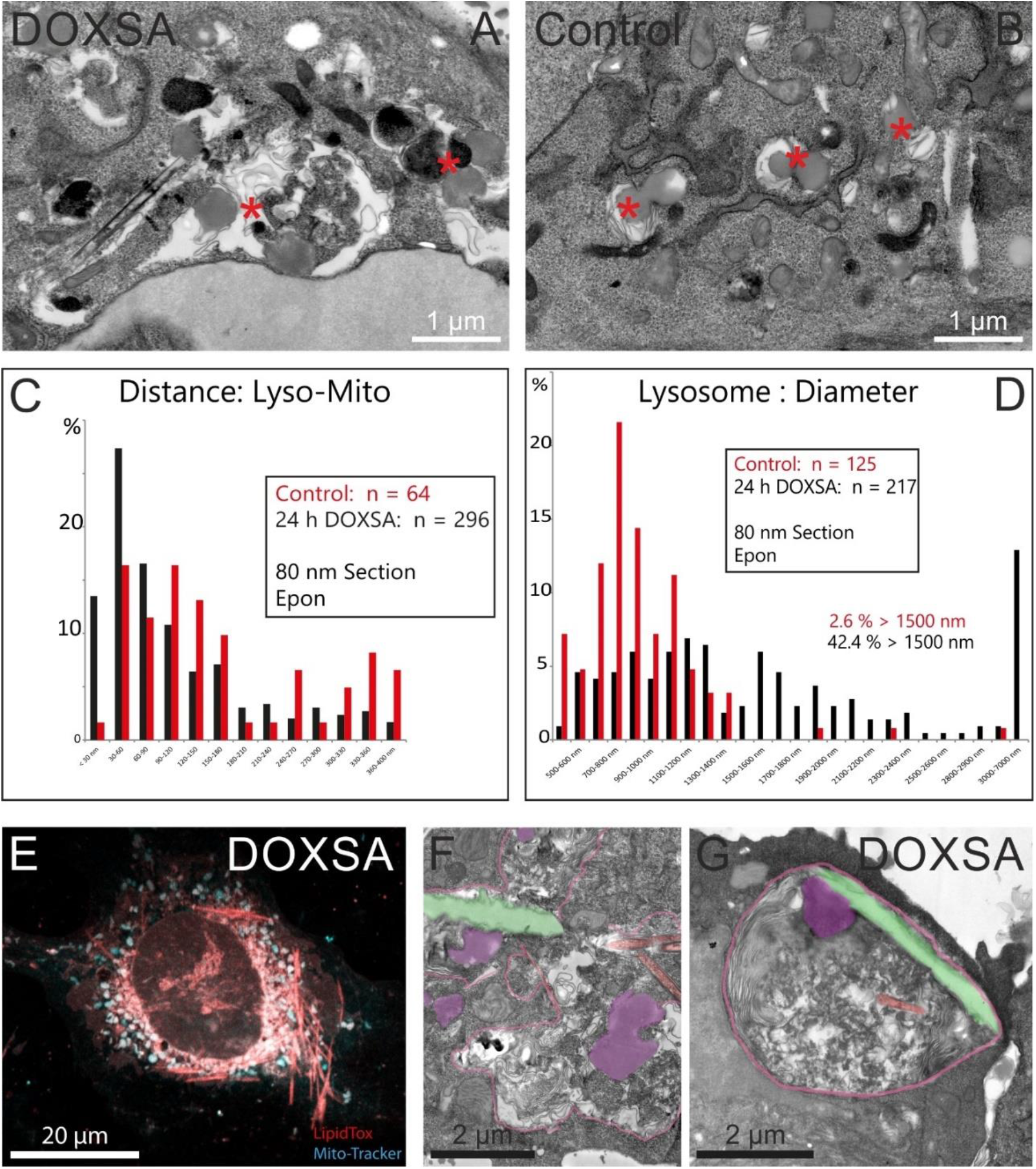
DeoxySL-mediated autophagy led to lysosomal swelling. A) EM of MEF cells revealed expanded auto-lysosomal compartments (pink asterisk) comprising lipid membrane bulks, lipid particles and cytoplasmic remnants after applying DOXSA. B) Lysosomes (pink asterisk) in control cells appeared round and small. Quantitative EM indicated C) increased contact of mitochondria with lysosomes and corroborated D) deoxySL-induced lysosomal expansion. E) LipidTox dye (red) stained large needle-shaped lipid containing inclusions in cells after applying DOXSA. F) + G) EM of corresponding structures revealed crystal-like lysosomal inclusions (green) in expanded lysosomes (rose).

Using EM we recognized also bundles of lipid crystals > 15 µm (Fig. 10 A, green). To understand the dynamics of emerging lysosomal crystal inclusions, we performed TL-imaging using acidophilic LysoTracker Red DND-99 (LysoTracker), which highlighted also crystal-like inclusions (example indicated in pink) (Fig. 9 B). Lysosomal inclusions emerged on average 4 h after applying DOXSA to cell culture medium. We recognized an initial growth phase of about 40 min followed by a stationary phase (Fig. 9 C, Video S4 + S5). To reveal interplay of deoxySLs with dynamic lipid crystals we applied 900 nM DOXSA together with 100 nM alkyne-DOXSA to MEF cells followed by GCM labeling. GCM suggested alkyne-positive lipid particles (magenta) interacting with lipid inclusions (green) in lysosomal compartments (red). Additionally, GCM validated alkyne-deoxySL content (presumably hydrophobic alkyne-deoxyCers and alkyne-deoxy(DH)Cers) in lipid crystals suggesting deoxySLs involved in lysosomal substrate accumulation (Fig. 10 D).

**Figure 10:**
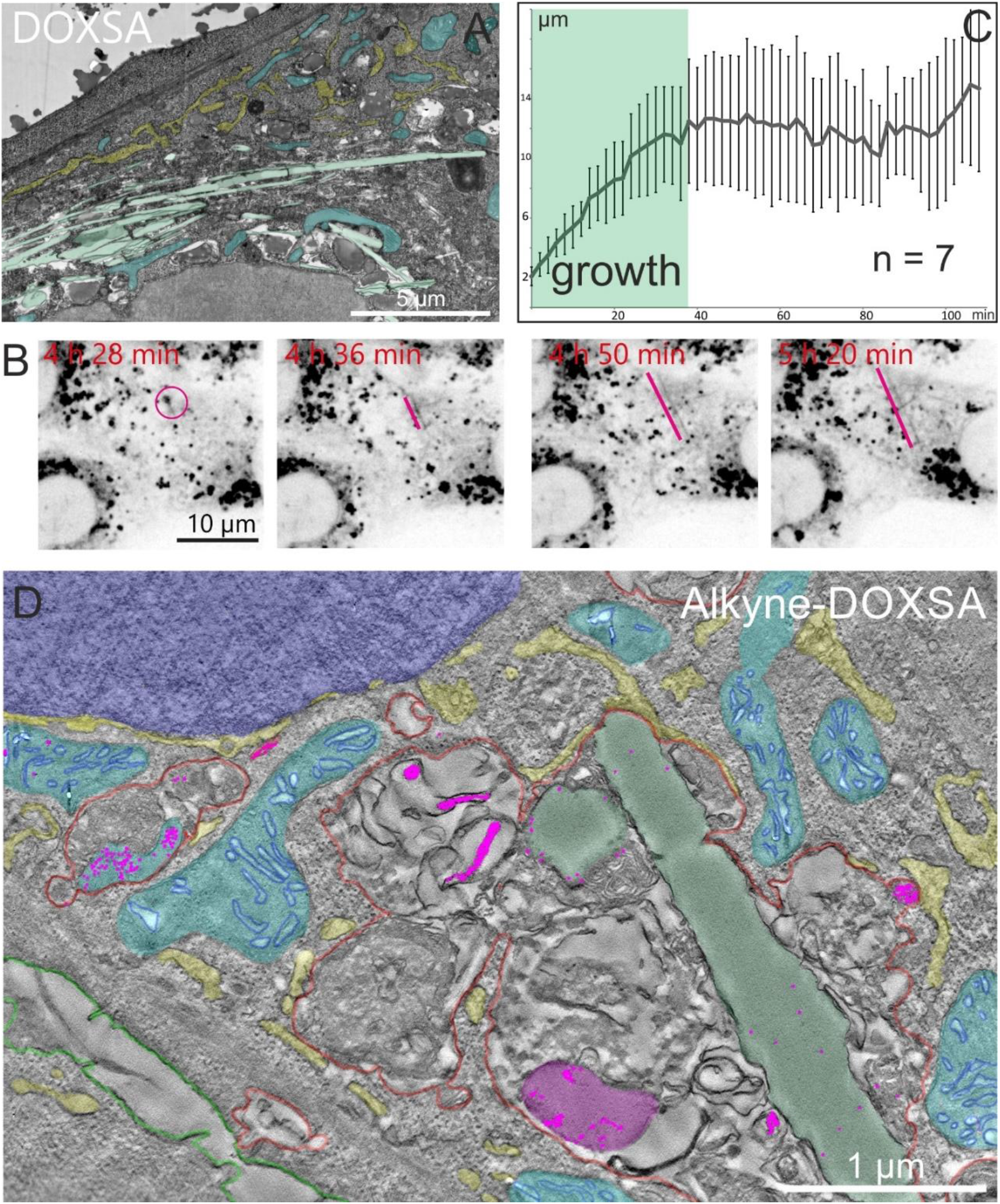
Applying DOXSA results in growth of crystal-like lysosomal inclusions. A) After applying 1 µM DOXSA to MEF cells, EM revealed lysosomal inclusions reaching a size > 15 µm (green). B) TL-imaging using LysoTracker demonstrated rapid growth of crystal-like lysosomal inclusions (indicated in pink) starting about 4 h after DOXSA-application. C) Quantifying observed dynamics revealed an initial growth phase of about 40 min followed by a stationary phase. D) GCM confirmed alkyne-deoxySL labeling (magenta dots) inside lipid-rich crystal-like lysosomal inclusions (green). (mitochondria = cyan; ER = yellow; lysosomal crystalloid = green, lipid particle = magenta; nucleus = blue, lysosome = red; GCM labeling = pink dots)

### DeoxySLs Tempted Lysosomal Exocytosis

Slow metabolic degradation of cytotoxic deoxySLs in a range of several days was proposed ^21 31^. However, mechanisms involved in detoxing cells after applying DOXSA remained obscure. EM of MEF cells treaded with 1 µM DOXSA revealed expanded lysosomes in the perinuclear area filled with dense cellular debris and organelle remnants (Fig. 10 A). GCM corroborated deoxySLs associated to lipid particles (magenta) and putative remnants of lipid crystals (green) inside lysosomes (red) (Fig 10 B). Additionally, EM revealed a population of lysosomes (red) fusing with the plasma membrane (green) suggesting lysosomal exocytosis of deoxySLs (Fig. 10 C). Lysosomal exocytosis of deoxySLs was substantiated by GCM revealing alkyne-positive lipid particles (magenta) in the extracellular space (Fig. 10 D). Indicated extracellular lipid particles (magenta) were confirmed using SEM of whole mount cell preparations of DOXSA-treated MEF cells and were largely missing in control samples.

## DISCUSSION

Increased intracellular DeoxySL levels have been shown to exert cytotoxic effects, while the underlying mechanisms of DOXSA and related cellular clearance processes remain still obscure. Previously intra-cellular accumulating deoxySLs were connected to cellular energy homeostasis^26^. Accordingly, strongly energy demanding neurons are sensitive against elevated DOXSA-levels and proliferative cell types exhibit an anti-proliferative effect after exogenously applying DOXSA. MEF cells turned out as a robust cell model for systematically examining DOXSA-related cell biological effects ^21 22 31^. Mitochondrial dysfunction, ER-stress, autophagy and accumulating lysosomes appeared connected to increased deoxySL-levels. In our hands deoxySL-mediated effects were restricted to replicating MEF cells, which is in accordance with observations of previous studies ^37^.

We applied 1 µM DOXSA to MEF cells, this amount was matching to the deoxySL-blood plasm level in patients suffering from diabetic neuropathies, HSNA1 and related tissue damage ^38 21^. Effects in cell culture resulted in decreased cell viability and appeared robust compared to prevalent DeoxySL-related clinical symptoms^57^. Distinct cytotoxic effects in single cells can be explained by the lack of lipid-buffering systems in the simplified in vitro arrangement. For tracing intra-cellular deoxySLs on ultra-structural level, we replaced 10 % of applied DOXSA with its traceable analogue alkyne-DOXSA as described previously^22^. In our study, examining lipid containing fine-structures in MEF cells was completed with TL-imaging of selected organelle tracers revealing DOXSA-mediated membrane dynamics.

Accurate spatial and metabolic tracing especially of bioactive lipids on ultra-structural level is of emerging interest to the scientific community. In previous approaches, biotin-labeled lipid analogues were employed to analyze the ultra-structural distribution of GM1 (monosialotetrahexosylganglioside), an SL involved into neurodegenerative GM1 gangliosidosis, along the endocytic route using gold-conjugated anti-biotin antibodies^58^. Another approach visualized propargyl-labeled phospholipid molecules, click-reacted with biotin-azide, by immuno-electron microscopy ^59^. Furthermore, azido-choline was provided to cells and its incorporation into cellular lipids was visualized using EM after click-reaction and diaminobenzidine (DAB) photo-oxidation^47^. Alkyne-DOXSA, enabling lipid tracing and quantitative analysis of lipid metabolism, was introduced and validated before ^22^. Here, we complement click-LM ^44^ with a new labeling approach for tracing alkyne-lipids in ultra-structurally preserved cellular microenvironments; the Golden Click Method (GCM) provides a one-step labeling approach for tracing metabolically tagged alkyne-lipids on the nanometer scale using EM. Amphiphilic ultra-small PEG1000-azide gold probes are exceptionally small and readily diffuse into chemically fixed cells without using detergents. Abandonment of detergents from sample preparation improved preservation of delicate lipid landmarks. After copper-catalyzed click-reaction in aldehyde fixed cells, an electrochemical silver deposition was used for enhancing and immobilizing ultra-small gold labeling followed by sample processing for EM modified from Karreman et al. 2014 ^60^. Cacodylate buffer (CB) appeared incompatible with GCM and was therefore replaced with HEPES resulting in a minor loss of sample contrast and preservation.

Based on predicted biophysical properties, an impact of local accumulating deoxyCers / deoxy(DH)Cers on sub-cellular lipid membranes is likely^25^. Accordingly, we and others showed DOXSA-mediated changes in mitochondrial ultra-structure with a loss of cristae before^22 61^. Additionally, forming of mitochondrial tubes containing deoxySLs was described ^31^. Here, we confirmed build-up of a deoxySL-fraction in extended lipid inclusions inside of tubular mitochondria. On ultra-structural level a clear-cut localization of alkyne-deoxySLs to the lumen of described lipid inclusions was evident. We hypothesize that the presence of a considerable quantity of poorly miscible deoxyCers / deoxy(DH)Cers in mitochondrial sub-compartments resulted in observed organelle distortion and loss of physiological functions.

Macroautophagy (here referred to as autophagy), a process characterized by double membraned structure (the autophagosome), is a key trafficking pathway required to transfer of damaged or obsolete cellular compartments to lysosomes for subsequent enzymatic degradation^62^. The closed autophagosome develops from an open membrane structure termed the isolation-membrane, which engulfs cytoplasmic constituents. Previous studies proposed the ER as lipid source for the isolation-membrane^63 64^. Subsequent fusion between the autophagosome and lysosomal compartments forms the autolysosome, where enclosed materials are degraded^65^. Cellular stress can result in mitochondrial dysfunction and lead subsequently to mitochondrial fission. While intact fragmented mitochondria can restore the reticular mitochondrial network, defective compartments, which lost its membrane potential, induce protective mitophagy ^66 67^. During mitophagy mitochondria are sequestered by an autophagosome and subsequently degraded by lysosomal fusion^68 69^. Additionally, ER-stress triggers autophagy and the ER-compartment can become a target of the degrading machinery. This process is referred as ER-phagy^70^. On ultra-structural level ER-phagy is characterized by vesicles densely filled with lipid membranes and little cytosol indicating that ER-membranes were sequestrated selectively ^71^. Previous studies indicated that exposure of MEF cells and neurons to DOXSA tempted ER-stress and an intoxication leading to mitochondrial fission and autophagy ^22 31^. However, detailed interplay between mitophagy, ER-phagy and deoxySLs was not described.

In our study, engulfment of tubular mitochondria due to intercalating membranes was not evident suggesting that mitophagy was not targeting extended dysfunctional mitochondria. However, tubular mitochondria interacted with lysosomes containing lipid. Observed mitochondria-lysosome interaction indicated metabolite transfer between both compartments. We also observed close contact between lysosomes and extended smooth ER-domains exhibiting putative deoxyCer / deoxy(DH)Cer containing lipid inclusions. We propose that detoxing extended mitochondria and ER from accumulating bioactive deoxyCers / deoxy(DH)Cers is a lysosome dependent process supporting cellular clearance and survival.

A parallel occurring consequence after applying DOXSA is mitochondrial fission and forming of hydrophobic bodies ^22 72 73^. We observed that fragmented mitochondria showing impaired cristae structure and accumulating deoxySLs were targeted by mitophagy. In contrast, mitochondria with intact cristae appeared not associated with deoxySLs and were less frequently targeted by degrading processes. We suggest that mitochondrial fission and mitophagy separate deoxySL-containing dysfunctional from functional mitochondrial sub-compartments preventing further organelle damage. A similar process appeared related to ER-phagy. Resulting substantial loss of intact mitochondria and ER leads to reduced cellular ATP-levels and impaired protein synthesis respectively^30^. The resulting cellular energy deficit can explain decreased cell proliferation after exogenously applying DOXSA. DeoxySL-mediated mitophagy and ER-phagy complements previous findings that deoxySLs increased the autophagic flux in MEF cells ^31^. Accordingly we observed that induced autophagy resulted eventually in an accumulating fraction of deoxyCers / deoxy(DH)Cers in lysosomes.

Complex lipids are degraded within lysosomes by soluble hydrolytic enzymes. If a lipid substrate cannot get degraded by the given set of lysosomal enzymes, it accumulates in the lysosome and can cause, together with coprecipitation of other hydrophobic substances, cellular injury. The resulting condition is comparable to a lysosomal “traffic jam” compromising lysosomal function. This effect can ultimately lead to cellular starvation^74^. DeoxyCers lack the hydroxyl group at the C1 preventing conversion of these metabolite to hydrophilic deoxySL-species; this prevents their degradation by sphingosine-1-phosphate lyase^21^. The resulting lipid storage leads to lysosomal dysfunction resulting in cellular starvation^74^. Accordingly, we observed expanded lysosomes comprising putative deoxyCer / deoxy(DH)Cer containing lipid particles interacting with rigid lipid crystals. We suggest that lipid crystals, which were previously related to inflammation processes, displayed a storage compartment for hydrophobic deoxySL substrates. However, the density of GCM labeling in intra-lysosomal lipid crystals was comparably low, indicating that alkyne-containing gold probes were hindered entering dense lipid crystals, which were previously indicated as birefringent^31^.

Additionally, we observed degraded cytoplasm and organelle remnants accumulated in lysosomes pointing to starvation induced non-selective autophagy^56^. Accordingly, lysosome-associated membranes sequestrating cytoplasm were evident. In contrast to conventional autophagosomes, the observed membrane arrangements were tubular and extended. Despite ultra-structural features resembling isolation-membranes, autophagosomes and a putative ER-origin, a final conclusion about the origin and composition of the described membrane compartments cannot be drawn here. DeoxySL-induced changes in cellular lipid composition and destabilizing effects on the cytoskeleton may represent possible explanations for extended intracellular membrane compartments.

An important factor that determines the development of lysosomal storage in cells is connected to strategies for disposing the stored material ^75^. Lysosomes can translocate to the plasma membrane for subsequent fusion resulting in release of their contents ^76^. This process, called lysosomal exocytosis, is an appealing option to dispose accumulated lysosomal substrates promoting cellular clearance after lysosomal substrate accumulation ^77^. Accordingly, we recognized a population of lysosomes loaded with lipid particles fused with the plasma membrane. Release of lysosomal content including putative deoxyCer / deoxy(DH)Cer containing lipid particles into the extracellular space was evident. We propose that this process contributed to cellular clearance of excess deoxySLs resulting in reactivation of cell proliferation within several days.

Our findings link cytotoxic deoxySLs, lipid membrane alterations and elementary cell biological processes, such as mitophagy, ER-phagy and abnormal lysosomal storage (Fig. 12). These insights may support new conclusions about tissue damage related to drug induced neuropathy, diabetes type 2 and HSNA1. Based on our data, future studies can confirm comparable structural effects in neurons and human tissue biopsies. In this context, our work may provide new diagnostic markers of clinically relevant symptoms and molecular targets suitable for medical intervention.

**Figure 11:**
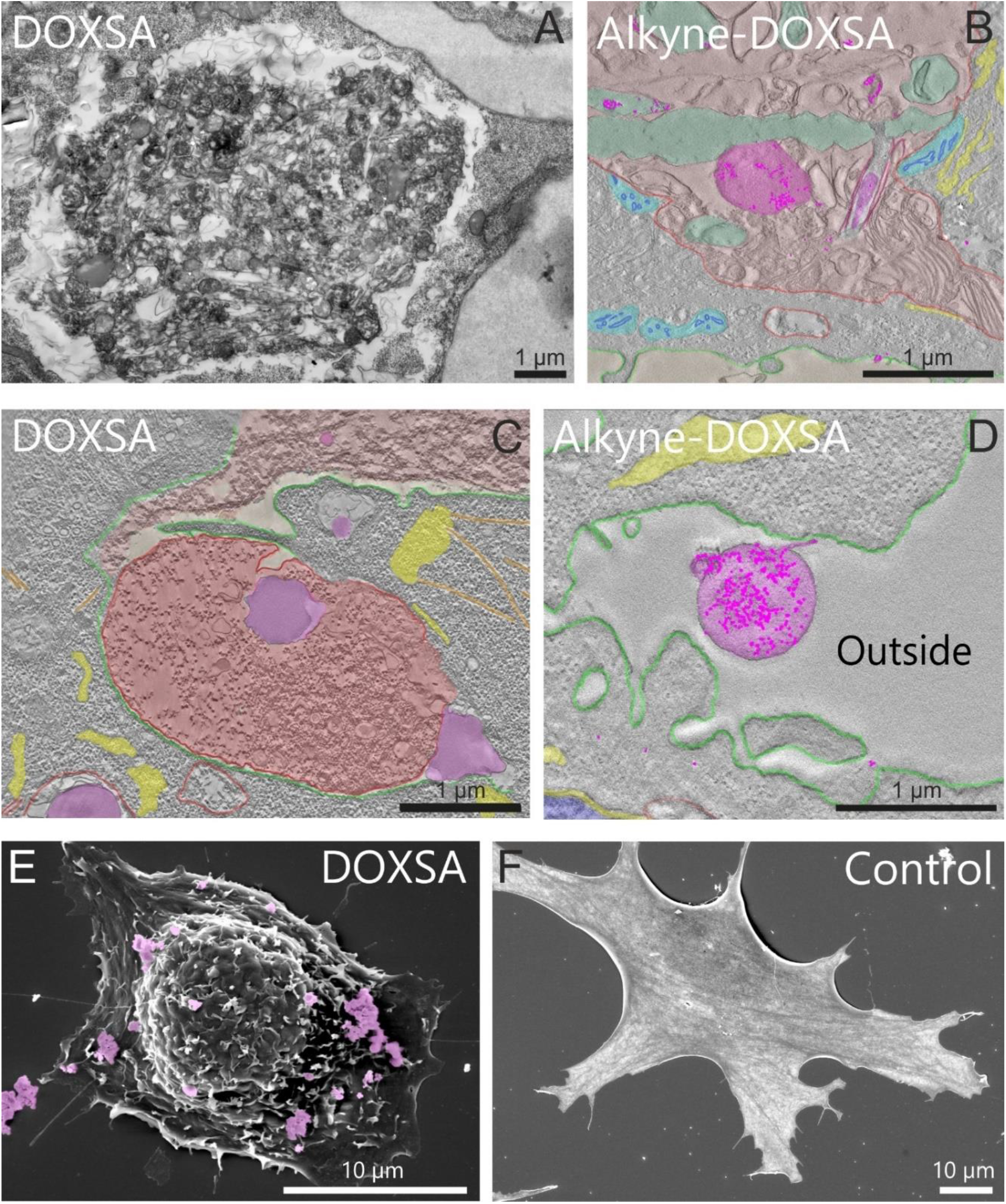
Lysosomal exocytosis of deoxySLs. A) EM indicated accumulating organelle remnants in lysosomes after applying 1 µm DOXSA to MEF cells. B) After applying 900 nM DOXSA and 100 nM alkyne-DOXSA, GCM confirmed DeoxySLs content in intra-lysosomal lipid particles (magenta) and putative remnants of crystal-like lysosomal inclusions (green). C) A population of lysosomes (red) was recruited to the plasma membrane (green) corroborating lysosomal exocytosis. D) Lysosomal exocytosis of deoxySLs was substantiated by GCM revealing alkyne-positive lipid particles (magenta) in the extracellular space. E + F) SEM of DOXSA treated MEF cells confirmed extracellular lipid particles (magenta). (mitochondria = cyan; ER: Yellow; lysosomal crystalloid [remnant] = green; lysosome = red; GCM labeling = pink dots).

**Figure 12:**
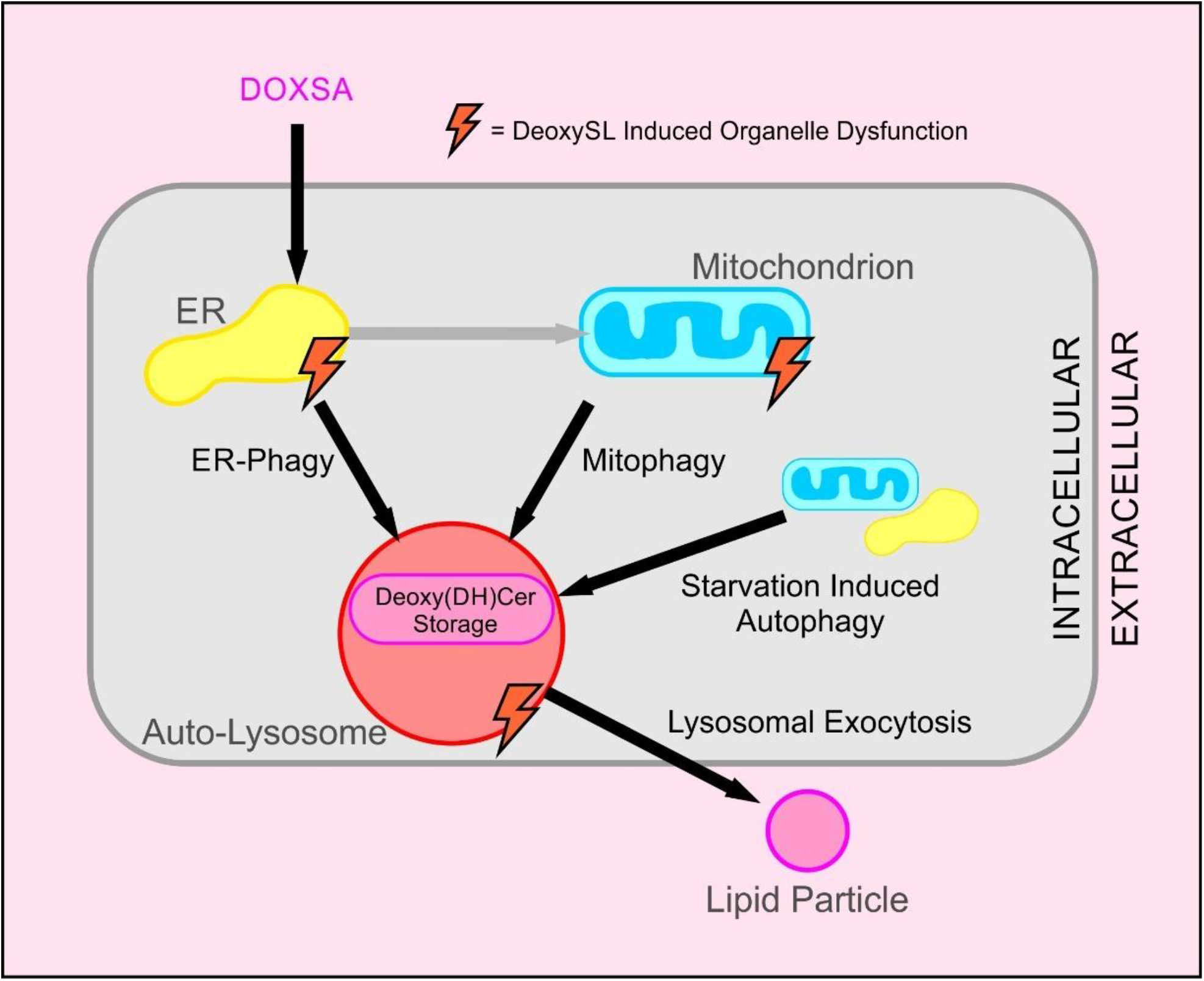
Model of 1-deoxy-sphingolipid mediated organelle dysfunction and subsequent cellular clearance of accumulated lysosomal lipid substrates. After applying DOXSA to cell culture medium, deoxySLs translocated to ER and mitochondria. ER-stress, mitochondrial dysfunction and a transient anti-proliferative effect were evident. ER-phagy and mitophagy of dysfunctional membrane compartments resulted in translocating slowly degradable deoxy(DH)Cers to auto-lysosomal compartments. Subsequent abnormal lysosomal lipid storage compromised lysosomal function. Multiple deoxySL-related organelle dysfunctions led to cellular starvation and connected non-selective autophagy. Lysosomal exocytosis of putative deoxy(DH)Cer containing lipid particles contributed to cellular clearance of cytotoxic deoxySLs promoting reactivation of cell proliferation within several days.

## Supporting information

Video S1 Time-lapse imaging of mitochondrial fission after applying DOXSA to MEF cells

Video S2 Live-imaging of an individual mitochondrial fission events after applying DOXSA to MEF cells.

Video S3 Time-lapse imaging of ER-membrane dynamics after applying DOXSA to MEF cells.

Video S4 Time-lapse imaging of lysosome dynamics after applying DOXSA to MEF cells

Video S5 Time-lapse imaging of crystal-like lysosomal lipid inclusions (Zoom in from Video S4)

## ACKNOWLEDGEMENTS

The authors thank Dr. H. U. Fried, Dr. I. König and Dr. C. Möhl (DZNE Bonn, Germany) for support regarding light microscopy and image processing. Also, thanks to Drs. S. Irsen and J. Feydt (Research Center Caesar, Bonn, Germany) and the “Electron Microscopy and Analytics” (EMA) team for assistance regarding electron microscopy. A grateful thanks to Drs. A. Al-Amoudi (DZNE Bonn, Germany), L. Kuerschner and C. Thiele for financial support and Dr. D. O. Fuerst (University of Bonn, Germany) for helpful scientific discussions. This research was funded by the Deutsche Forschungsgemeinschaft (SFB-TRR83 to C.L. and M.H.)

## SUPPLEMENT

**Video S1: Time-lapse imaging of mitochondrial fission after applying DOXSA to MEF cells**. A 5 h TL-acquisition (1 frame / 1 min) using spinning disc microscopy after adding 100 nM MitoTracker Green FM confirmed disintegration of the mitochondrial network after applying 1 µM DOXSA (right) compared to vehicle control (left). (Scale bar = 5 µm)

**Video S2: Live-imaging of an individual mitochondrial fission events after applying DOXSA to MEF cells**. Live-imaging (5 frames / 1 sec) using spinning disc microscopy after adding 100 nM MitoTracker Green FM demonstrated individual DOXSA-mediated mitochondrial fission events. (Scale bar = 2 µm)

**Video S3: Time-lapse imaging of ER-membrane dynamics after applying DOXSA to MEF cells**. A 12.5 h TL-acquisition (1 frame / 5 min) using spinning disc microscopy after adding 100 nM ER-Tracker Blue-White DPX revealed ER-membrane dynamics after applying 1 µM DOXSA (right) compared to vehicle control (left). (Scale bar = 5 µm)

**Video S4: Time-lapse imaging of lysosome dynamics after applying DOXSA to MEF cells**. A 8 h TL-acquisition (1 frame / 2 min) using spinning disc microscopy after adding 50 nM Lystracker Red DND-99 revealed lysosmal dynamics after applying 1 µM DOXSA (right) compared to vehicle control (left). (Scale bar = 5 µm)

**Video S5: Time-lapse imaging of crystal-like lysosomal lipid inclusions (Zoom in from Video S4)**. A TL-acquisition (1 frame / 2 min) using spinning disc microscopy after applying 50 nM Lystracker Red DND-99 demonstrated rapid growth of crystal-like lysosomal inclusions. (Scale bar = 5 µm)

**Fig. S1:**
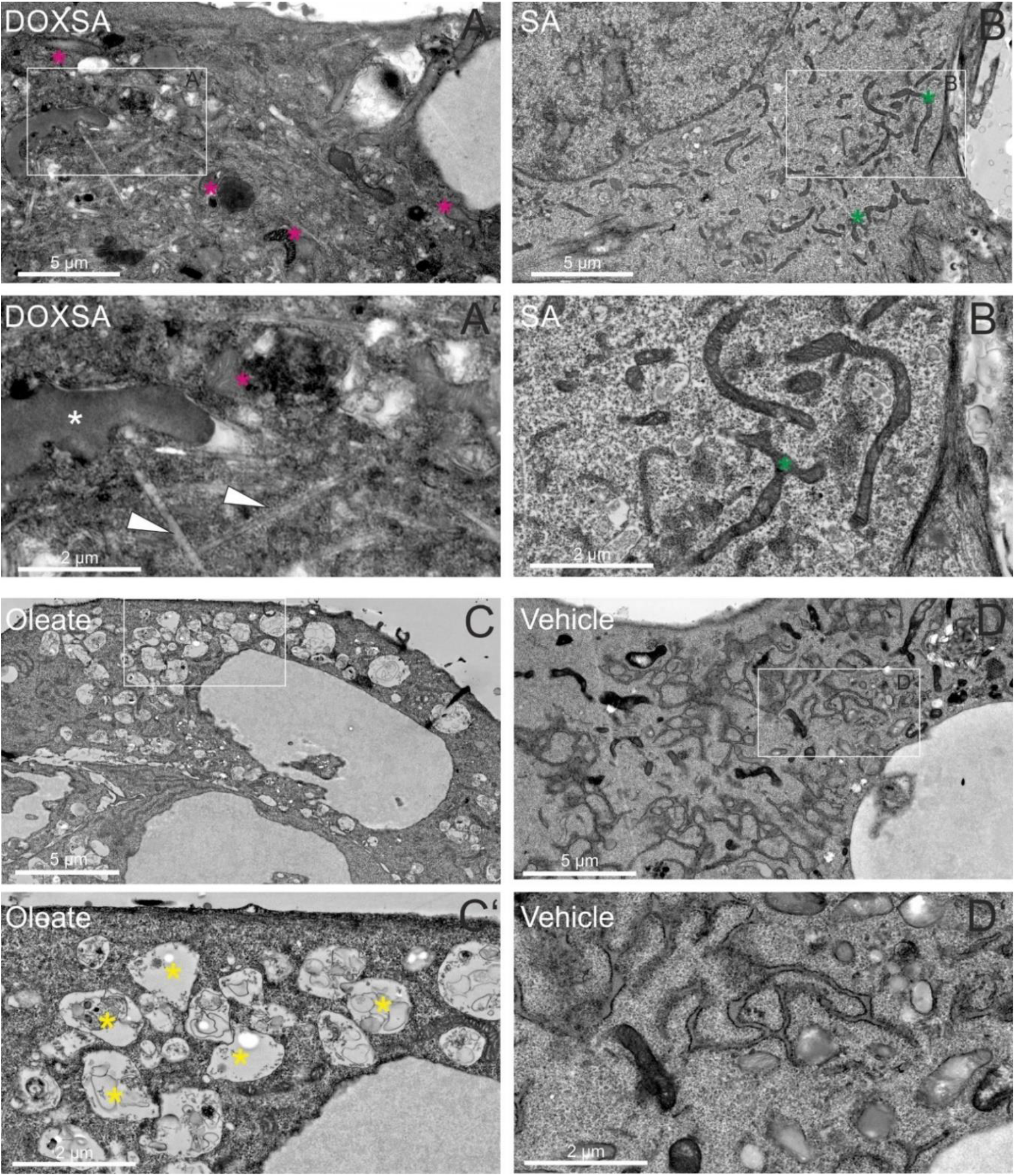
Applying DOXSA tempts characteristic lipid accumulation and organelle distortion in MEF cells. MEF cells were treated with A) 1 µm DOXSA, B) 1 µM sphinganine (SA), C) 20 µM Oleate (Ole) or D) vehicle control (0.1 % Ethanol) and examined using EM. A’,B’C’ and D’) show framed areas in A – D). Applying 1 µM DOXSA for 24 h tempted extended (arrowheads) and spherical (white asterisk) lipid inclusions and revealed fragmented mitochondria (pink asterisk). In contrast treatment with 1 µM SA for 24 h indicated hyper-fused mitochondria (green asterisk). Applying 20 µM Ole for 24 h resulted in swollen lysosomes (yellow asterisk), while vehicle control appeared unaffected.

**Fig. S2:**
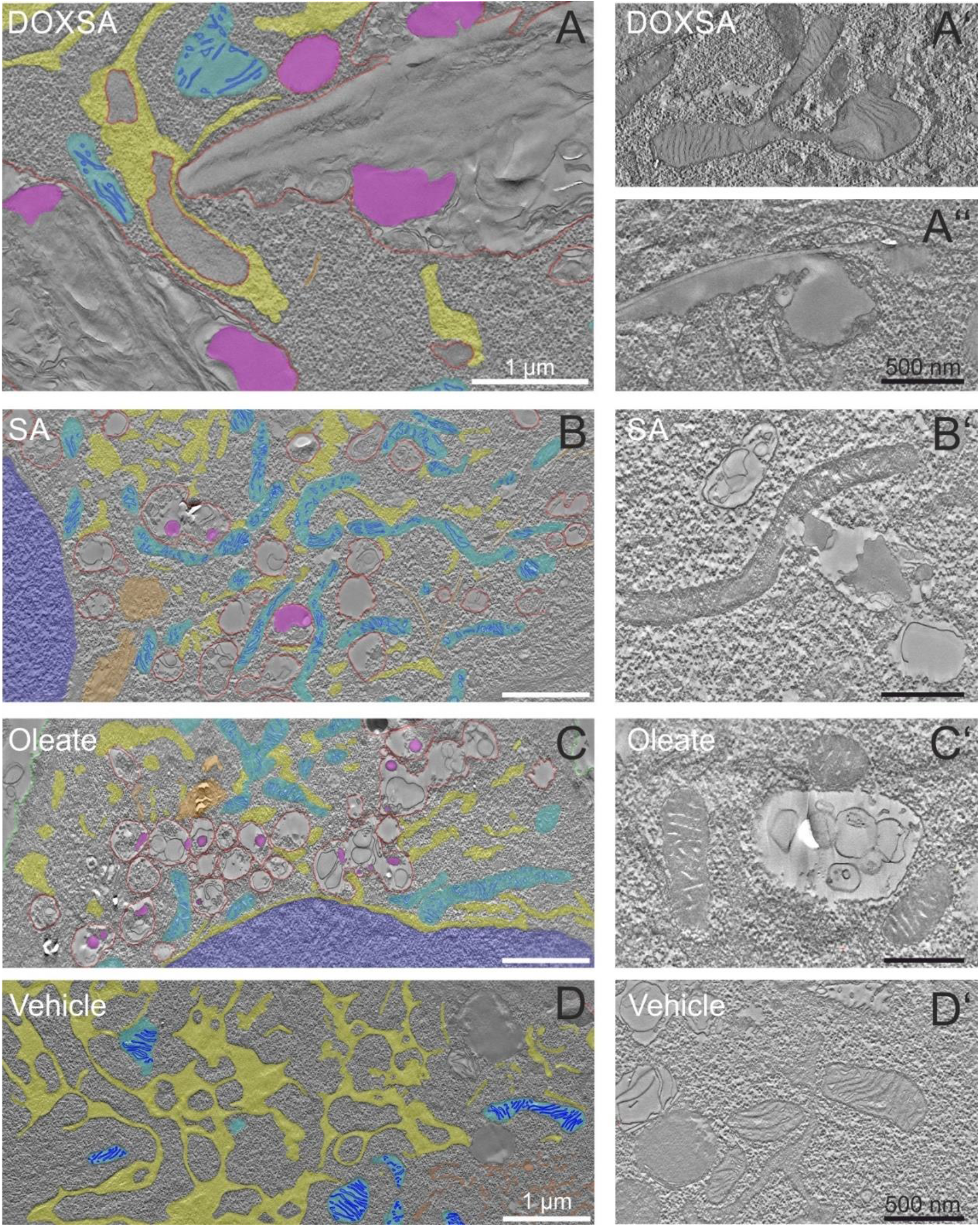
Applying DOXSA tempts characteristic lipid accumulation, ER-collapse and mitochondrial fission in MEF cells.: MEF cells were treated with A) 1 µm DOXSA, B) 1 µM sphinganine (SA), C) 20 µM Oleate (Ole) or D) vehicle control (0.1 % Ethanol) and examined using ET. A’, A’, B’, C’ and D’) show representative reconstructions without overlay. While DOXSA-treated cells were characterized by lipid aggregates (magenta), fragmented mitochondria (cyan), collapsed ER (yellow) and expanded lysosomes (red), ET showed smaller lysosomes (red) interacting with hyper-fused mitochondria (cyan) after applying SA. Lysosomes (red) induced by 20 µM Oleate were characterized by small lipid inclusions (magenta) and appeared also closely related to mitochondria (cyan).

**Fig. S3:**
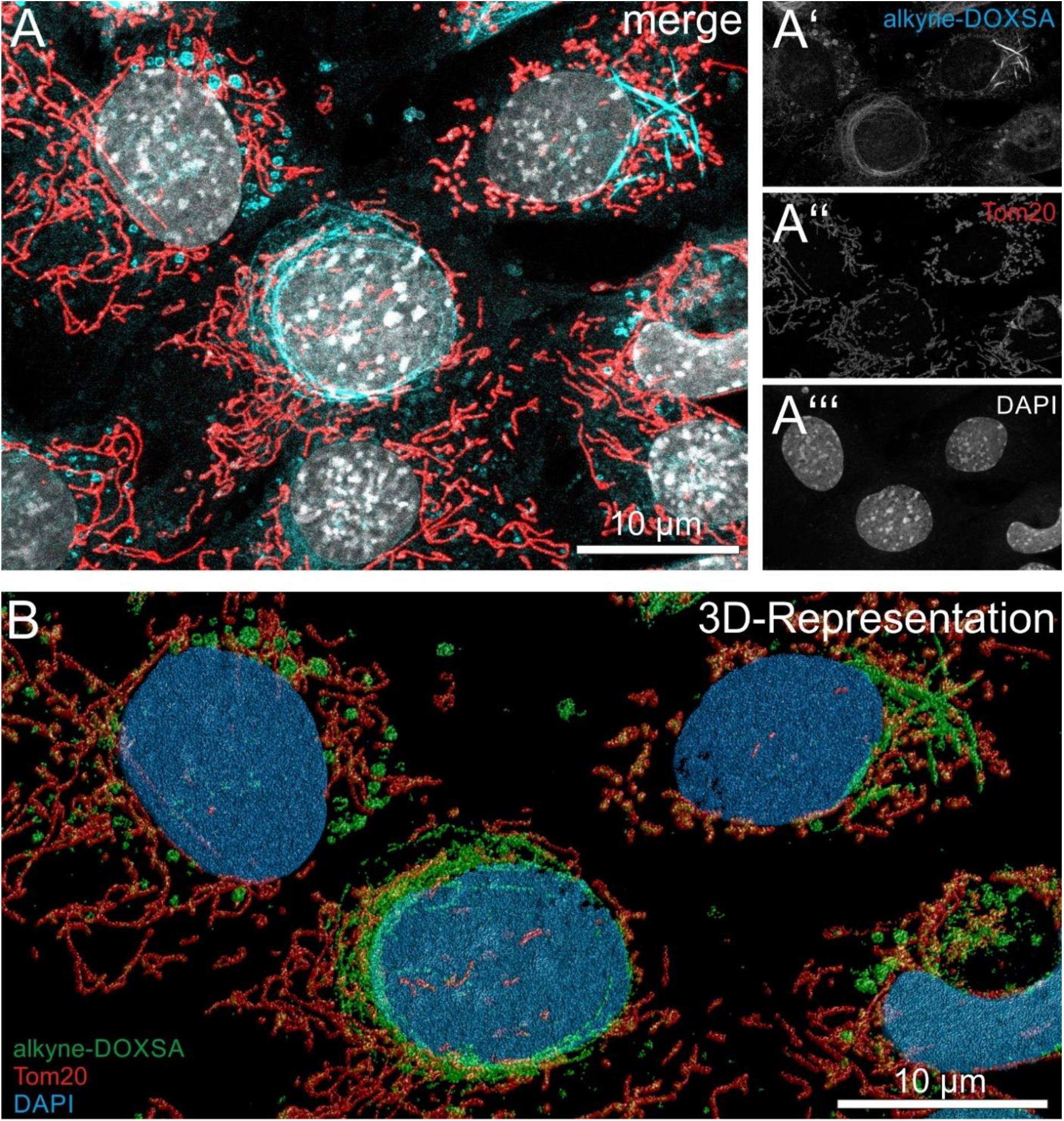
LSM reveled alkyne-deoxySL positive structures in MEF cells. MEF cells were treated with 900 nM DOXSA and 100 nM alkyne-DOXSA for 24 h. A) After fixing cells, co-labeling for alkyne-deoxySLs (cyan) and the mitochondrial marker Tom20 (red) was performed together with DAPI-staining (blue). Alkyne-deoxySLs were click-reacted with ASTM-BODIPY; the secondary antibody for Tom20 labeling was conjugated with Alexa 555. Micrographs were recorded using laser scanning microscopy (LSM) before deconvolution using Huygens software. A maximum intensity projection (MIP) revealed alkyne-deoxySL labeling in spherical structures, in mitochondria and prominently in different extended compartments. A’-A’’’) depicts separate channels of A). B) A 3D-Representation of the deconvolved image stack, generated using Huygens software, represents the intra-cellular alkyne-lipid distribution (green) in the whole volume of selected MEF cells (Tom20 = red, DAPI = blue). LM revealed diverse sub-cellular patterns of alkyne-lipid labeling in adjacent MEF cells representing different stages of DOXSA-intoxication.

**Fig. S4:**
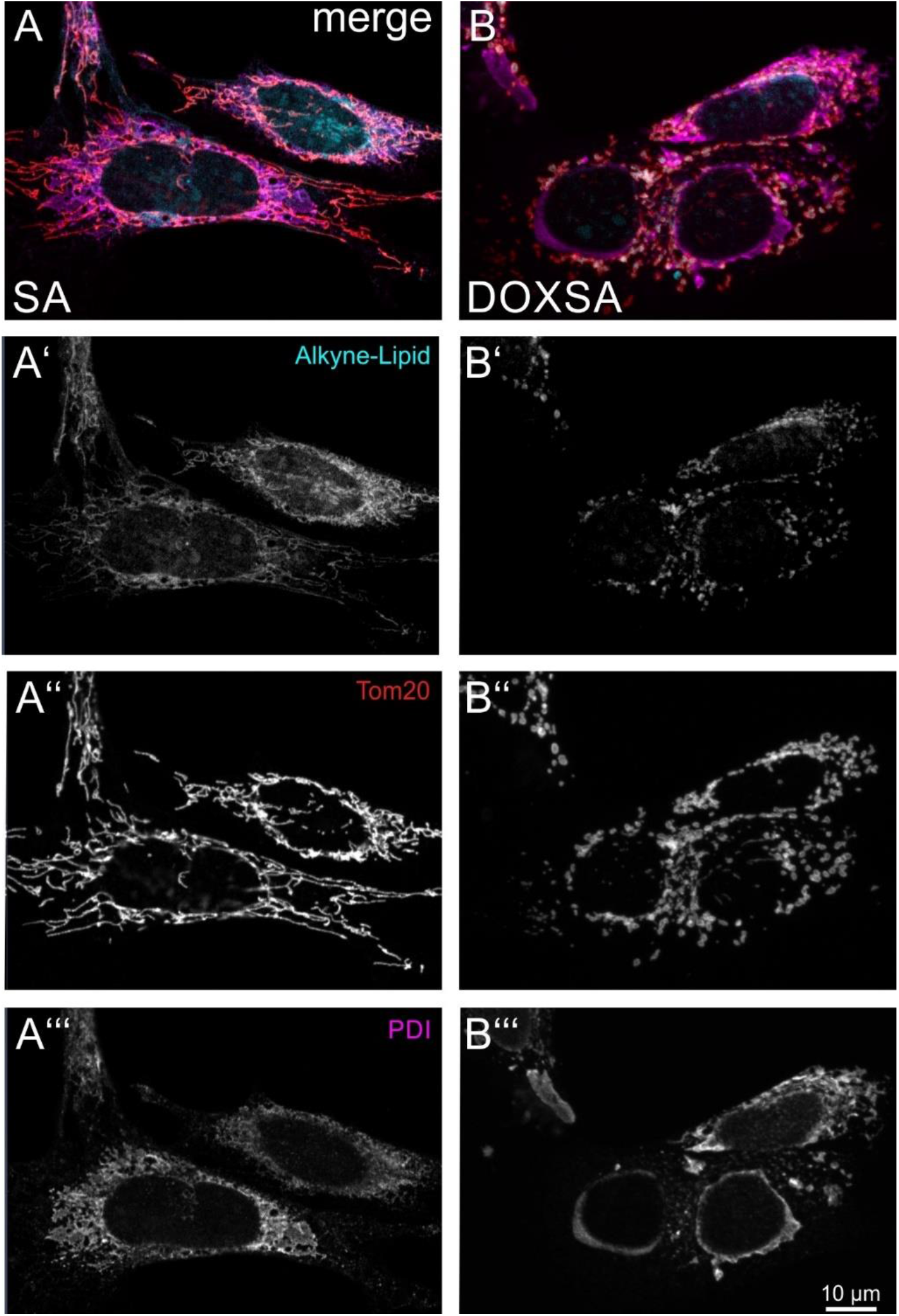
DeoxySLs induced mitochondrial fission and ER-collapse. A +B) MEF cells, treated with 900 nM SA and 100 nM alkyne-SA or 900 nM DOXSA and 100 nM alkyne-DOXSA for 24 h, were compared after co-labeling for alkyne-deoxySLs (cyan), the mitochondrial marker Tom20 (red) and the rough ER-marker PDI (magenta). Alkyne-lipids were click-reacted with ASTM-BODIPY; the secondary antibody for PDI labeling was conjugated with Alexa 555 and the secondary antibody for Tom20 labeling with Alexa 647. Imaging was done using LSM. A’-A’’’) and B’-B’’’) depict separate channels of A) and B). Mitochondrial fission and ER-collapse was only observed after applying DOXSA. In depicted cells alkyne-DOXSA localized to fragmented mitochondria, there was no evidence for co-localization of alkyne-labeling with PDI. Alkyne-SA labeling co-localized typically with the reticular mitochondrial network.

